# Optimum growth temperature declines with body size within fish species

**DOI:** 10.1101/2021.01.21.427580

**Authors:** Max Lindmark, Jan Ohlberger, Anna Gårdmark

**Affiliations:** Swedish University of Agricultural Sciences, Department of Aquatic Resources, Institute of Coastal Research, Skolgatan 6, Öregrund 742 42, Sweden; School of Aquatic and Fishery Sciences (SAFS), University of Washington, Box 355020, Seattle, WA 98195-5020, USA; Swedish University of Agricultural Sciences, Department of Aquatic Resources, Skolgatan 6, SE-742 42 Öregrund, Sweden

**Keywords:** body growth, metabolic rate, consumption rate, temperature-size rule, metabolic theory of ecology, climate change, warming, scaling, allometry

## Abstract

According to the temperature-size rule, warming of aquatic ecosystems is generally predicted to increase individual growth rates but reduce asymptotic body sizes of ectotherms. However, we lack a comprehensive understanding of how growth and key processes affecting it, such as consumption and metabolism, depend on both temperature and body mass within species. This limits our ability to inform growth models, link experimental data to observed growth patterns, and advance mechanistic food web models. To examine the combined effects of body size and temperature on individual growth, as well as the link between maximum consumption, metabolism and body growth, we conducted a systematic review and compiled experimental data on fishes from 52 studies that combined body mass and temperature treatments. By fitting hierarchical models accounting for variation between species, we estimated how maximum consumption and metabolic rate scale jointly with temperature and body mass within species. We found that whole-organism maximum consumption increases more slowly with body mass than metabolism, and is unimodal over the full temperature range, which leads to the prediction that optimum growth temperatures decline with body size. Using an independent dataset, we confirmed this negative relationship between optimum growth temperature and body size. Small individuals of a given population may therefore exhibit increased growth with initial warming, whereas larger conspecifics could be the first to experience negative impacts of warming on growth. These findings help advance mechanistic models of individual growth and food web dynamics and improve our understanding of how climate warming affects the growth and size structure of aquatic ectotherms.

## Introduction

Individual body growth is a fundamental process powered by metabolism, and thus depends on body size and temperature (Brown et al., 2004). It affects individual fitness and life history traits, such as maturation size, population growth rates (Savage et al., 2004), and ultimately energy transfer across trophic levels (Andersen et al., 2009; Barneche & Allen, 2018). Therefore, understanding how growth scales with body size and temperature is important for predicting the impacts of global warming on the structure and functioning of ecosystems.

Global warming is predicted to lead to declining body sizes of organisms (Daufresne et al., 2009; Gardner et al., 2011). The temperature size-rule (‘TSR’) states that warmer rearing temperatures lead to faster developmental times (and larger initial size-at-age or size-at-life-stage), but smaller adult body sizes in ectotherms (Atkinson, 1994; Ohlberger, 2013). This relationship is found in numerous experimental studies (Atkinson, 1994), is reflected in latitudinal gradients (Horne et al., 2015), and is stronger in aquatic than terrestrial organisms (Forster et al., 2012; Horne et al., 2015). Support for the TSR exists in fishes, in particular in young fish, where reconstructed individual growth histories often reveal positive correlations between growth rates and temperature in natural systems (Baudron et al., 2014; Huss et al., 2019; Neuheimer et al., 2011; Thresher et al., 2007). However, whether the positive effect of warming on growth is indeed limited to small individuals within a species, as predicted by the temperature size-rule, is less clear. Negative correlations between maximum size, asymptotic size or size-at-age of old fish and temperature have been found in commercially exploited fish species (Baudron et al., 2014; Ikpewe et al., 2020; van Rijn et al., 2017). However, other studies, including large scale experiments, controlled experiments and latitudinal studies or observational data on unexploited species, have found no or less clear negative relationships between maximum size, growth of old fish or mean size and temperature (Audzijonyte et al., 2020; Barneche et al., 2019; Denderen et al., 2020; Huss et al., 2019; van Dorst et al., 2019) and differences between species may be related to life history traits and depend on local environmental conditions (Denderen et al., 2020; Wang et al., 2020).

While the support for TSR is mixed, and the underlying mechanisms are not well understood (Audzijonyte et al., 2019; Neubauer & Andersen, 2019; Ohlberger, 2013), theoretical growth models, such as Pütter growth models (Pütter, 1920), including the von Bertalanffy growth model (VBGM) (von Bertalanffy, 1957), commonly predict declines in asymptotic body mass with temperature and declines in optimum growth temperature with body mass, in line with the TSR (Morita et al., 2010; Pauly, 2021; Pauly & Cheung, 2018a; Perrin, 1995). Yet, the physiological basis of these models has been questioned, as the commonly applied scaling parameters (mass exponents) tend to differ from empirical estimates (Lefevre et al., 2018; Marshall & White, 2019). Hence, despite attempting to describe growth from first principles, Pütter growth models can also be viewed as phenomenological. In more mechanistic growth models, the difference between energy gain and expenditure is partitioned between somatic growth and gonads (Essington et al., 2001; Jobling, 1997; Kitchell et al., 1977; Ursin, 1967). Energy gain is normally the amount of energy extracted from consumed food, and expenditure is defined as maintenance, activity and feeding metabolism. These components of the energetics of growth are found in dynamic energy budget models (Kitchell et al., 1977; Kooijman, 1993), including those used in physiologically structured population models (PSPMs) (de Roos & Persson, 2001) and size-spectrum models (Blanchard et al., 2017; Hartvig et al., 2011; Maury & Poggiale, 2013). Therefore, it is important to understand how consumption and metabolism rates scale with body mass and temperature in order to understand if and how body growth of large fish within populations is limited by temperature, and to evaluate the physiological basis of growth models.

Moreover, the effect of body mass and temperature on growth dynamics should be evaluated over ontogeny at the intraspecific level (within species), which better represents the underlying process than interspecific data (among species) (Marshall & White, 2019). For instance, we are not aware of an interspecific relationship between optimum growth temperature and body mass, but within species it may have a large effect on growth dynamics. Despite this, intraspecific body mass and temperature scaling is often inferred from interspecific data, and we know surprisingly little about average relationship between consumption and metabolic exponents within species (Marshall & White, 2019). Importantly, how physiological rates depend on mass and temperature within species can differ from the same relationships across species (Glazier, 2005; Jerde et al., 2019; Rall et al., 2012). Across species, rates are often assumed and found to scale as power functions of mass with exponents of 3/4 for whole organism rates, exponentially with temperature, and with independent mass and temperature effects (e.g., in the Arrhenius fractal supply model (AFS) applied in the metabolic theory of ecology, MTE (Brown et al., 2004; Downs et al., 2008; Gillooly et al., 2001)). In contrast, within species, deviations from a general 3/4 mass exponent are common (Barneche et al., 2019; Bokma, 2004; Clarke & Johnston, 1999; Jerde et al., 2019), rates are typically unimodally related to temperature, activation energies can vary a lot (Dell et al., 2011; Englund et al., 2011; Pawar et al., 2016; Rall et al., 2012; Uiterwaal & DeLong, 2020) and the effects of mass and temperature can be interactive (García García et al., 2011; Glazier, 2005; Lindmark et al., 2018; Ohlberger et al., 2012; Xie & Sun, 1990) (but see Jerde *et al*. (2019)). Extensions of the MTE include fitting multiple regression models where coefficients for mass and temperature are estimated jointly (Downs et al., 2008), as well as fitting non-linear models that can capture the de-activation of biological rates at higher temperatures (Dell et al., 2011; Englund et al., 2011; Padfield et al., 2017; Schoolfield et al., 1981). To advance our understanding of the intraspecific properties of mass- and temperature dependence of biological rates, intraspecific data with variation in both mass and temperature are needed.

In this study, we analyzed how maximum consumption and metabolic rate of fish scale intraspecifically with mass and temperature, and how optimum growth temperature scales with size. We performed a systematic literature review by searching the Web of Science Core Collection to compile datasets on individual-level maximum consumption, metabolic and growth rate of fish from experiments in which the effect of fish body mass is replicated across multiple temperatures within species (total n=3672, with data from 20, 34 and 13 species for each rate, respectively). We then fit hierarchical Bayesian models to estimate general intraspecific scaling parameters while accounting for variation between species. The estimated mass dependence and temperature sensitivity of consumption and metabolism were used to quantify average changes in net energy gain (and hence, growth, assumed proportional to net energy gain) over temperature and body mass. Lastly, we compared our predicted changes in optimum growth temperature over body mass with an independent experimental dataset on optimum growth temperatures data across individuals of different sizes within species.

## Materials and methods

### Data acquisition

We searched the literature for experimental studies evaluating the temperature response of individual maximum consumption rate (feeding rate at unlimited food supply, *ad libitum*), resting, routine and standard oxygen consumption rate as a proxy for metabolic rate (Nelson, 2016) and growth rates across individuals of different sizes within species. We used three different searches on the Web of Science Core Collection (see *SI Appendix*, for details). In order to estimate how these rates depend on body size and temperature within species, we selected studies that experimentally varied both body size and temperature (at least two temperature treatments and at least two body masses). The average number of unique temperature treatments (temperature rounded to nearest °C) by species is 7.2 for growth and 4.3 for consumption (below peak temperatures) and metabolism data. The criteria for both mass and temperature variation in the experiments reduce the number of potential data sets, as most experimental studies use either size or temperature treatments, not both. However, these criteria allow us to fit multiple regression models and estimate the effects of mass and temperature jointly, and to evaluate the probability of interactive mass- and temperature effects within species. Following common practice we excluded larval studies, which represents a life stage exhibiting different constraints and scaling relationships than non-larval life stages (Glazier, 2005).

Studies were included if (i) a unique experimental temperature was recorded for each trial (±1°C), (ii) fish were provided food at *ad libitum* (consumption and growth data) or if they were unfed (resting, standard or routine metabolic rate), and (iii) fish exhibited normal behavior during the experiments. We used only one study per species and rate to ensure that all data within a given species are comparable as measurements of these rates can vary between studies due to e.g. measurement bias, differences in experimental protocols, or because different populations were studied (Armstrong & Hawkins, 2008; Jerde et al., 2019). In cases where we found more than one study for a given rate and species, we selected the most suitable study based on our pre-defined criteria (for details, see *SI Appendix*). We ensured that the experiments were conducted at ecologically relevant temperatures (*SI Appendix,* Figs. S1, S3). A more detailed description of the search protocol, data selection, acquisition, quality control, collation of additional information and standardizing of rates to common units can be found in *SI Appendix*.

We compiled four datasets: maximum consumption rate, metabolic rate, growth rate and the optimum growth temperature for each combination of body mass group and species. We compiled a total of 746 measurements of maximum consumption rate (of which 666 are below peak), 2699 measurements of metabolic rate and 227 measurements of growth rate (159 below peak, 45 optimum temperatures for species-size group combinations) from published articles for each rate, from 20, 34 and 13 species, respectively, from different taxonomic groups, habitats and lifestyles (Table S1-S2). We requested original data from all corresponding authors of each article. In cases where we did not hear from the corresponding author, we extracted data from tables or figures using Web Plot Digitizer (Rohatgi, 2012).

### Model fitting

#### Model description

To each dataset, we fit hierarchical models with different combinations of species-varying coefficients, meaning they are estimated with shrinkage. This reduces the influence of outliers which could occur in species with small samples sizes (Gelman & Hill, 2007; Harrison et al., 2018). The general form of the model is:

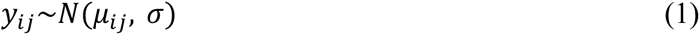

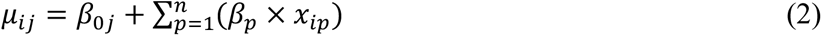

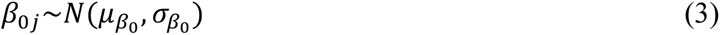

where *y*_ij_ is the *i*th observation for species *j* for rate *y*, *β*_0j_ is a species-varying intercept, *x*_i*p*_ is a predictor and *β*_*p*_ is its coefficient, with *p* = 1, . . . , *n*, where *n* is the number of predictors considered in the model (mass, temperature, and their interaction). Predictors are mean centered to improve interpretability (Schielzeth, 2010). Species-level intercepts follow a normal distribution with hyperparameters μ_*β*_0__ (global intercept) and σ_*β*_0__ (between-species standard deviation). For most models we also allow the coefficient *β*_*p*_ to vary between species, such that *β*_*p*_ becomes *β*_*p*j_ and *x*_i*p*_ *x*_ij*p*_, where *β*_*p*j_∼*N*(μ_*βp*_, σ_*βp*_). For each dataset, we evaluate multiple combinations of species-varying coefficients (from varying intercept to *n* varying coefficients). We used a mix of flat, weakly informative, and non-informative priors. For the temperature and mass coefficients we used the predictions from the MTE as the means of the normal prior distributions (Brown et al., 2004), but with large standard deviations (see *SI Appendix*, Table S3). Below we describe how the model in Eqns. 1-3 is applied to each data set.

#### Mass- and temperature dependence of maximum consumption and metabolic rate below peak temperatures

Peak temperature (optimum in the case of growth) refers to the temperature at which the rate was maximized, by size group. For data below peak temperatures, we assumed that maximum consumption and metabolic rate scale allometrically (as a power function of the form *I* = *i*_0_*M*^*b*0^ ) with mass, and exponentially with temperature. Hence, after log-log (natural log) transformation of mass and the rate, and temperature in Arrhenius temperature (1/*kT* in unit eV^-1^, where *k* is Boltzmann’s constant [8.62×10^-5^ eV K^-1^]), the relationship between the rate and its predictors becomes linear. This is similar to the MTE, except that we estimate all coefficients instead of correcting rates, and allow intercepts and slopes to vary across species.

When applied to Eqns. 1-3, *y*_ij_ is the *i*th observation for species *j* of the natural log of the rate (maximum consumption or metabolic rate), and the predictors are *m*_ij_ (natural log of body mass), *t*_*A*,ij_ (Arrhenius temperature, 1/*kT* in unit eV^-1^), both of which were mean-centered, and their interaction. Body mass is in g, consumption rate in g day^-1^ and metabolic rate in mg O_2_ h^-1^. We use resting or routine metabolism (mean oxygen uptake of a resting unfed fish only showing some spontaneous activity) and standard metabolism (resting unfed and no activity, usually inferred from extrapolation or from low quantiles of routine metabolism, e.g. lowest 10% of measurements) to represent metabolic rate (Beamish, 1964; Ohlberger et al., 2007). Routine and resting metabolism constitute 58% of the data used and standard metabolism constitutes 42%. We accounted for potential differences between these types of metabolic rate measurements by adding two dummy coded variables, *type*_*r*_ and *type*_*s*_, the former taking the value 0 for standard and 1 for a routine or resting metabolic rate measurement, and vice versa for the latter variable. Thus, for metabolism, we replace the overall intercept *β*_0j_ in Eqns. 2-3 with *β*_0*r*j_ and *β*_0*s*j_. *β*_0*s*j_ is forced to 0 for a species that has a routine or resting metabolic rate and *β*_0*r*j_ is forced to 0 for a species with standard metabolic rate data. We assume these coefficients vary by species following normal distributions with global means μ_*β*0r_ and μ_*β*0*s*_, and standard deviations σ_*β*0r_ and σ_*β*0*s*_, i.e. *β*_0*r*j_∼*N*(μ_*β*0r_, σ_*β*0r_) and *β*_0*s*j_∼*N*(μ_*β*0*s*_, σ_*β*0*s*_).

#### Mass- and temperature dependence of consumption including beyond peak temperatures

Over a large temperature range, many biological rates are unimodal. We identified such tendencies in 10 out of 20 species in the consumption data set. To characterize the decline in consumption rate beyond peak temperature, we fit a mixed-effects version of the Sharpe Schoolfield equation (Schoolfield et al., 1981) as expressed in (Padfield et al., 2020), to equations 1-2 with *y*_ij_ as rescaled consumption rates (*C*). Specifically, we model μ_ij_ in Eq. 1 with the Sharpe-Schoolfield equation:

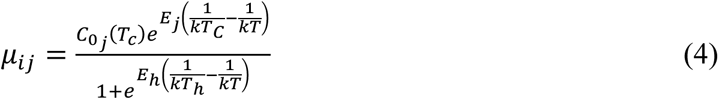

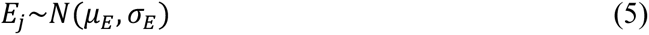

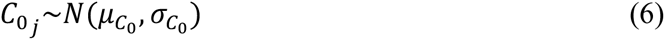

where *C*_0j_(*T*_*C*_) is the rate at a reference temperature *T*_*C*_ in Kelvin [K] (here set to 263.15, note it is on centered scale), *E*_j_ [eV] is the activation energy, *E*_7_ [eV] characterizes the decline in the rate past the peak temperature and *T*_7_ [K] is the temperature at which the rate is reduced to half (of the rate in the absence of deactivation) due to high temperatures. We assume *E*_j_ and *C*_0j_ vary across species according to a normal distribution with means μ_*E*_ and μ_*C*0_, and standard deviations σ_*E*_ and σ_*C*0_ (Eq. 5-6). Prior to rescaling maximum consumption (in unit g day^-1^) by dividing *C*_i,j_ with the mean within species 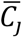, we isolate the effect of mass by dividing consumption with *m*^0.63^, which is the estimated allometric relationship from the log-linear model. Temperature, *T*, is centered by subtracting the temperature at peak consumption. This was estimated separately for each species using the Sharpe-Schoolfield equation but without group-varying coefficients in a frequentist framework (see ‘*Parameter estimation*’). The rescaling is done to control for differences in the optimum temperature between species, which if not accounted for obscures the average relationship between consumption and temperature among species.

#### Mass-dependence of optimum growth temperature

To evaluate how the optimum temperature (*t*_*opt*,ij_, in degrees Celsius) for individual growth depends on body mass, we fit Eqns. 1-3 with *y*_ij_ as the mean-centered optimum growth temperature within species 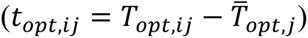, to account for species being adapted to different thermal regimes. *m*_ij_, the predictor variable for this model, is the natural log of the ratio between mass and mass at maturation acquired from FishBase (Froese & Pauly, 2019), within species: 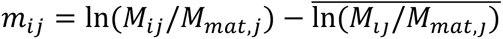. This rescaling is done because we are interested in examining relationships within species over “ontogenetic size”, and because we do not want to confound that effect with any relationship that might occur across species that have different asymptotic sizes. We consider both the intercept and the effect of mass to potentially vary between species.

#### Parameter estimation

We fit the models in a Bayesian framework, using R version 4.0.2 (R Core Team, 2020) and JAGS (Plummer, 2003) through the R-package ‘*rjags*’ (Plummer, 2019). We used 3 Markov chains with 5000 iterations for adaptation, followed by 15000 iterations burn-in and 15000 iterations sampling where every 5^th^ iteration saved. Model convergence was assessed by visually inspecting trace plots and potential scale reduction factors (Ȓ^T^) (*SI Appendix*). Ȓ compares chain variance with the pooled variance, and values <1.1 suggest all three chains converged to a common distribution (Gelman et al., 2003). We relied heavily on the R packages within ‘*tidyverse’* (Wickham et al., 2019) for data processing, as well as ‘*ggmcmc*’ (Fernández-i-Marín, 2016), ‘*mcmcviz*’ (Youngflesh, 2018) and ‘*bayesplot*’ (Gabry et al., 2019) for visualization. Frequentist single-species Sharpe-Schoolfield models were fitted using the packages ‘*rTPC’* (Padfield & O’Sullivan, 2020) and ‘*nls.multstart’* (Padfield & Matheson, 2020)

#### Model comparison

We compared the parsimony of models with different hierarchical structures, and with or without mass-temperature interactions, using the Watanabe-Akaike information criterion (WAIC) (Vehtari et al., 2017; Watanabe, 2013), which is based on the posterior predictive distribution. We report WAIC for each model descried above (Table S4-S5), and examine models with ΔWAIC values < 2, where ΔWAIC is each models difference to the lowest WAIC across models, in line with other studies (Olmos et al., 2019).

#### Net energy gain

The effect of temperature and mass dependence of maximum consumption and metabolism (proportional to biomass gain and losses, respectively) (Essington et al., 2001; Kitchell et al., 1977; Ursin, 1967) on growth is illustrated by visualizing the net energy gain. The model for the net energy gain (growth) can be viewed as an empirical temperature-dependent Pütter-type model. Pütter-type models are the simplest growth models based on a dynamic energy budget, and make strong assumptions about mass-scaling of key life-history and physiological processes (e.g., maturation and assimilation). However, Pütter-type models are among the most commonly applied growth models in ecology and fisheries, they tend to fit data reasonably well (Marshall & White, 2019), and are suitable for illustrating the consequences of non-linear consumption rates due to their simplicity (in contrast to more complex and parameter-rich dynamic energy budget models, e.g. (Cuenco et al., 1985; Kitchell et al., 1977)). A Pütter model is the result of two antagonistic allometric processes, biomass gains and biomass losses:

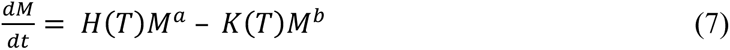

where *M* is body mass and *T* is temperature, *H* and *K* the allometric constants and *a* and *b* the exponents of the processes underlying gains and losses, respectively. We convert metabolism from oxygen consumption [mg O_2_ h^-1^ day^-1^] to g day^-1^ by assuming 1 kcal = 295 mg O_2_ (based on an oxycaloric coefficient of 14.2 J mg^-1^ O_2_) (Hepher, 1988), 1 kcal = 4184 J and an energy content of 5.6 kJ g^-1^ (wet weight) (Rijnsdorp & Ibelings, 1989). Hence, to convert our estimated intercept for routine/resting metabolic rates from unit log(mg O_2_ h^-1^ day^-1^) to g day^-1^, we multiply exp(μ_*β*_0__ ) with 0.0608063. Consumption rate is already in unit g day^-1^. Consumption and metabolic rates are calculated for two sizes (5 and 1000 g, which roughly correspond to the 25^th^ percentile of both datasets and the maximum mass in the consumption data, respectively), using the global allometric relationships found in the log-log models fit to sub-peak temperatures. These allometric functions are further scaled with the temperature correction factors *r*_*c*_ for consumption and *r*_*m*_ for metabolism. *r*_*c*_ is based on the Sharpe-Schoolfield model and *r*_*m*_ is given by the temperature dependence of metabolic rate from the log-linear model. Because *r*_*c*_ and *r*_*m*_ are fitted to data on different scales, we divide these functions by their maximum. Lastly, we rescale the product between the allometric functions and *r*_*c*_ and *r*_*m*_ such that the rate at 19°C (mean temperature in both data sets) equals the temperature-independent rate.

## Results

We identified that within species of fish, metabolic rates increase faster with body mass than maximum consumption rates, and neither of these rates conform to the commonly predicted 3/4 scaling with body mass (Fig. 1). We also quantified the unimodal relationship of consumption rate over the full temperature range (Fig. 2). Combined, these scaling relationships lead to the prediction, based on Pütter-type growth models, that optimum growth temperature declines with body size (Fig. 3). The prediction of declining optimum growth temperatures with size was confirmed by our analysis of independent experimental growth rate data. We find that within species the optimum growth temperature declines with body size by 0.31°C per unit increase in the natural log of relative body mass (Fig. 4). Below we present the underlying results in more detail.

**Figure 1.**
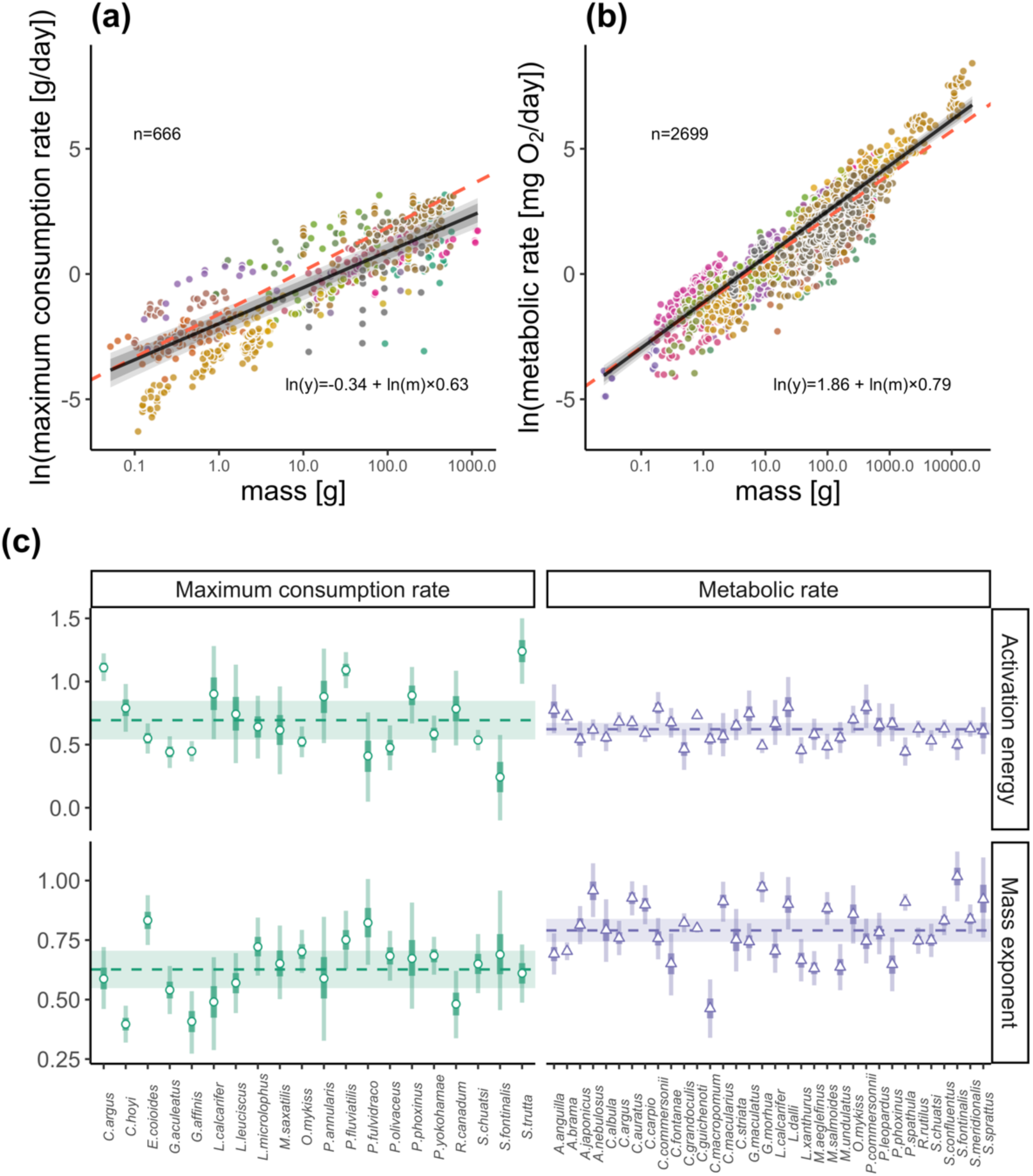
Natural log of maximum consumption rate (A) and metabolic rate (B) against body mass on a logarithmic x-axis. Lines and equation are global predictions on the centered data, x-axes are non-centered, (routine metabolic rate in panel B) at the average temperature in each data set (both 19°C, but note the model is fitted using mean-centered Arrhenius temperature), hence the temperature terms are omitted. Red dashed lines indicate a slope of 3/4, corresponding to the prediction from the metabolic theory of ecology. Gray bands correspond to 80% and 95% credible intervals. Species are grouped by color (legend not shown, n=20 for consumption and n=34 for metabolism, respectively). C) Global and species-level effects of mass and temperature on maximum consumption rate and metabolic rate. Horizontal lines show the posterior medians of the global mass exponents and activation energies of maximum consumption and metabolism (μ_*β*_0__ and −μ_*β*1_ (negative since we present activation energies rather than the coefficient) in Eq. 3 for the mass and temperature coefficients, respectively). The shaded horizontal rectangles correspond to the posterior median ± 1.96 standard deviations. Points and triangles show the posterior medians for each species-level coefficient (for maximum consumption rate and metabolic rate, respectively), and the vertical bars show their 80% and 95% credible interval.

**Figure 2.**
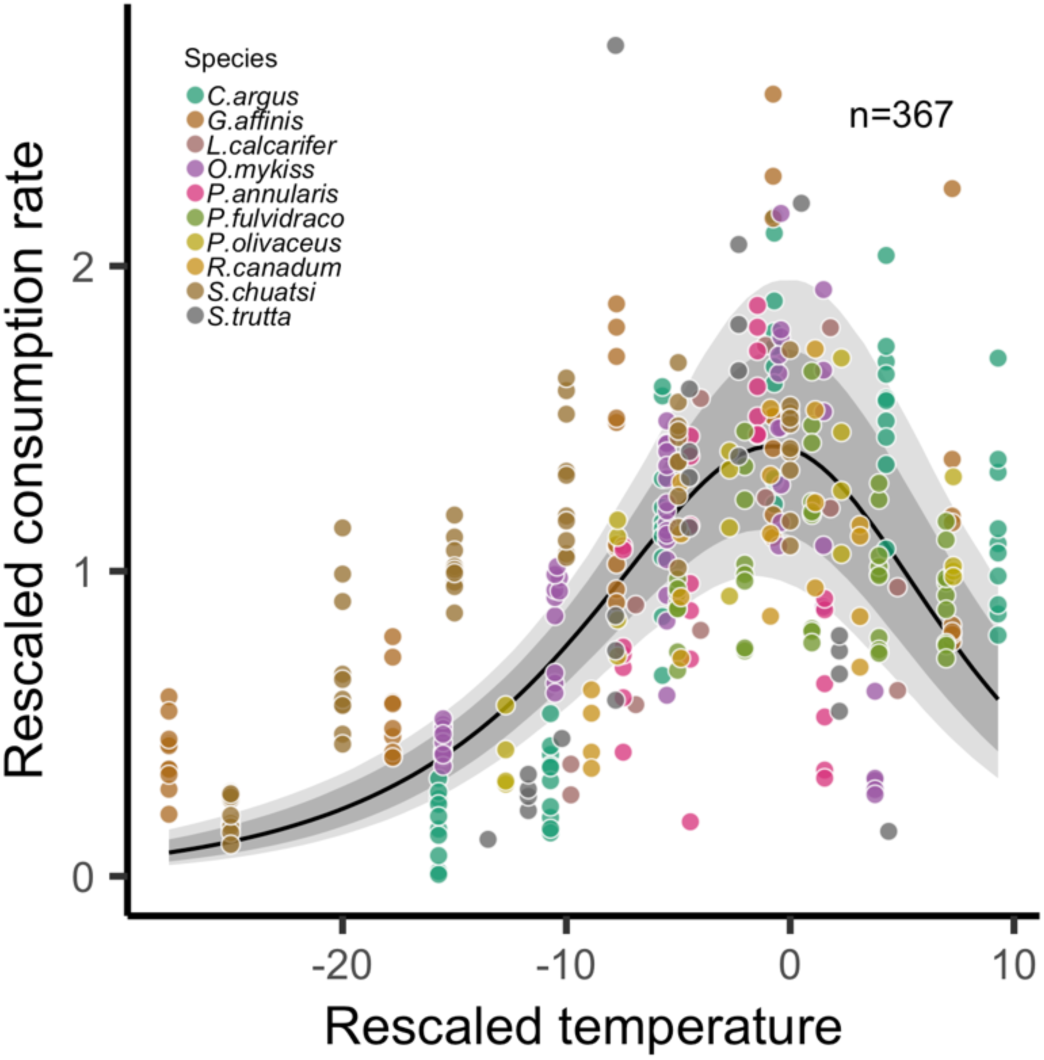
Mass-corrected maximum consumption rate increases until a maximum is reached, after which it declines steeper than the initial rate of increase. Maximum consumption rates are relative to the average maximum consumption rates within species and temperature is the difference between the experimental temperature and the temperature where maximum consumption peaks (also by species). The black line shows the posterior median of predictions from the Sharpe-Schoolfield model. Gray bands correspond to 80% and 95% credible intervals. Colors indicate species.

**Figure 3.**
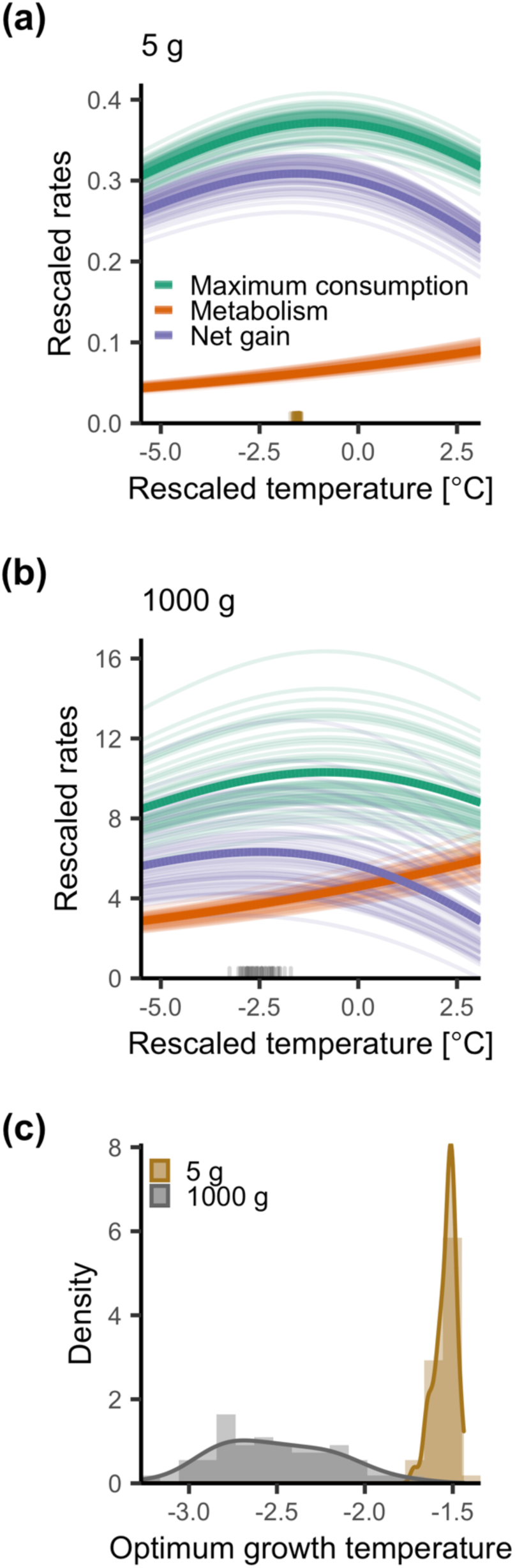
Illustration of predicted whole-organism maximum consumption rate (green), metabolic rate (orange) and the difference between them (‘net gain’; purple) for two body sizes (top=5g, center=1000g) over temperature (A-B) (see ‘Materials and Method’), and the distribution of optimum growth temperatures (C), i.e., the temperature at the peak of the net gain curve. Each line (50 in total) is a draw from the posterior distribution of the respective mass-scaling exponent, to illustrate uncertainty in the criteria leading to declining optimum growth temperatures with size. The thick solid lines correspond to the posterior median estimate (as reported in the ‘Result’).

**Figure 4.**
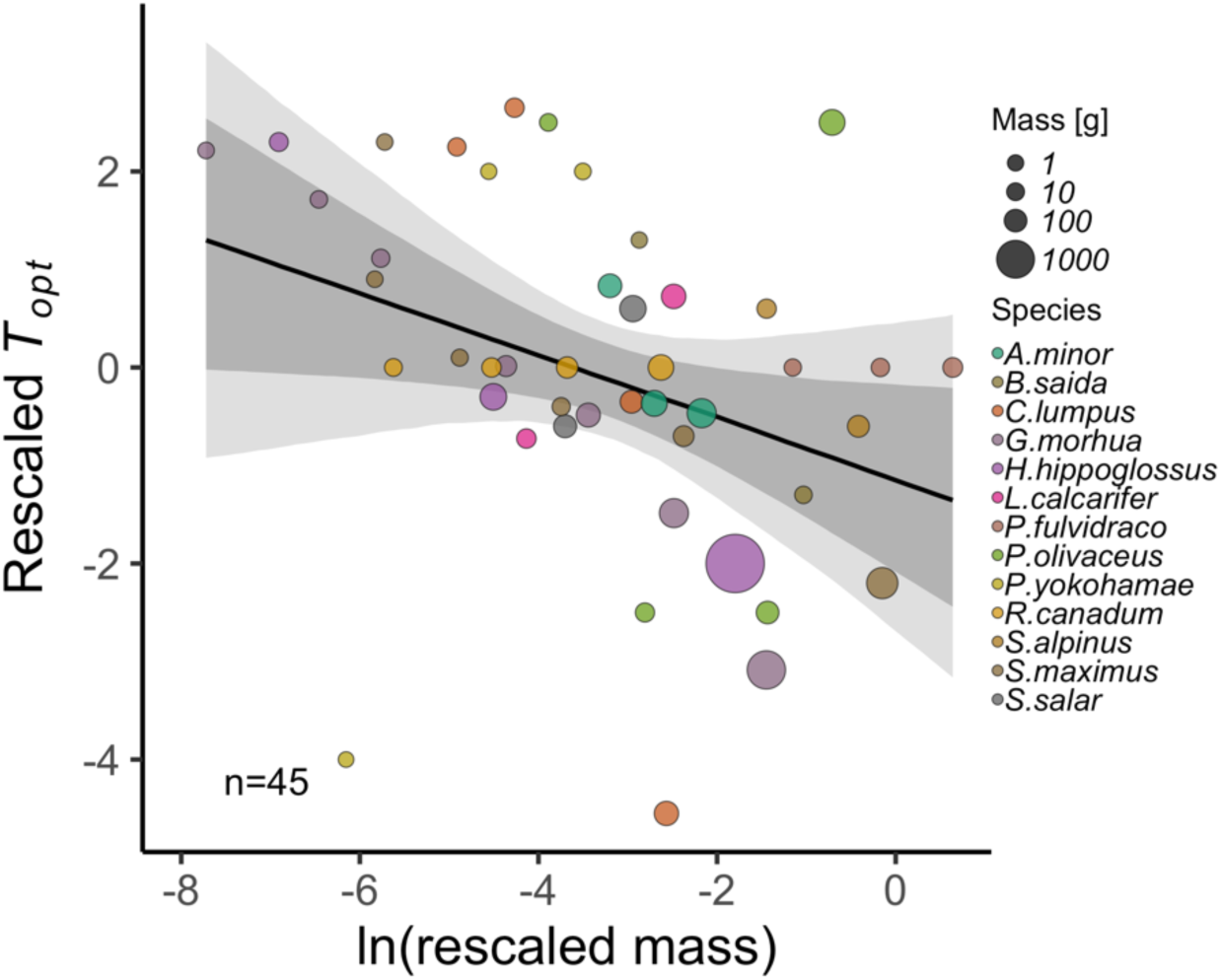
Experimental data demonstrating optimum growth temperature declines with intraspecific body mass. The plot shows the optimum temperature within species (rescaled by subtracting the mean optimum temperature from each observation, by species) as a function of the natural log of rescaled body mass (ratio of mass to maturation mass within species). The solid line shows the global prediction (using μ_*β*_0__ and μ_*β*1_ ), and shaded areas correspond to 80% and 95% credible intervals. Colors indicate species and the area of the circle corresponds to body mass in unit g.

We found that the average intraspecific mass exponent for consumption rate is smaller (0.63 [0.55, 0.71]) than that for metabolic rate (0.79 [0.74, 0.84]), based on the non-overlapping Bayesian 95% credible intervals (Fig. 1). It is also probable that the scaling exponents differ from 3/4 (that is predicted by the MTE), because >99% of the posterior distribution of the mass exponent of maximum consumption is below 3/4, and 95% of the posterior distribution of the mass exponent of metabolic rate is above 3/4. Activation energies of maximum consumption rate and metabolism are both similar (0.69 [0.54, 0.85] and 0.62 [0.57, 0.67] respectively; Fig. 1) and largely fall within the prediction from the MTE (0.6-0.7 eV) (Brown et al., 2004). The global intraspecific intercept for maximum consumption rate is estimated to be -0.34 [-0.85, 0.17], and for routine and resting metabolic rate it is estimated to be 1.85 [1.68, 2.04], and for standard metabolic rate it is 1.29 [0.97, 1.61] (*SI Appendix,* Fig. S6). Models where all coefficients varied by species were favored in terms of WAIC (M5 and M1, for consumption and metabolism, respectively) (*SI Appendix*, Table S4). We found statistical support for a species-varying mass and temperature interaction for metabolic rate; 98% of the posterior distribution of the global interaction coefficient μ_*β*_0__ is above 0 (*SI Appendix,* Fig. S4). The estimated coefficient is 0.0018 [0.015, 0.037] on the Arrhenius temperature scale, which corresponds to a decline in the mass scaling exponent of metabolic rate by 0.0026 °C^*1^. The selected model for maximum consumption rate did not include an interaction term between mass and temperature (M5).

We estimated the parameters of the Sharpe-Schoolfield equation (Eq. 4) for temperature-dependence of consumption including data beyond peak temperature as: activation energy, *E*_j_ = 0.73 [0.54, 0.94], rate at reference temperature, *C*_0j_ = 0.79 [0.58, 0.99], temperature at which the rate is reduced to half (of the rate in the absence of deactivation) due to high temperatures, *T_h_* = 0.75 [-0.86, 2.37], and the rate of the decline past the peak, *E_h_* = 1.89 [1.68, 2.1]. This shows that the relationship between consumption rate and temperature is unimodal and asymmetric, where the decline in consumption rate at high temperatures is steeper than the increase at low temperatures (Fig. 2).

The above results provide empirical support for the two criteria outlined in Morita et al., (2010) that result in declining optimum temperatures with size, i.e. (i) smaller whole organism mass exponent for consumption than metabolism (Fig. 1) and (ii) that growth reaches an optimum over temperature. In our case, the second criterion is met because consumption reaches a peak over temperature (Fig. 2) (in contrast to Morita *et al*. (2010), who assumed consumption to be linearly related to temperature, based on data from Atkinson (1994)). We illustrate the consequence of these findings in Fig. 3, which shows that the optimum temperature for net energy gain is reached at a lower temperature for a larger fish because of the difference in mass exponents of consumption and metabolism and because consumption is unimodally related to temperature. Assuming growth is proportional to net energy gain, this predicts that optimum growth temperature declines with body size.

Using independent data from growth trials across a range of body sizes and temperatures, we also find statistical support for a decline in optimum growth temperature with body mass within species, because 92% of the posterior density of the global slope estimate (μ_*β*1_) is below 0. The models with and without species-varying slopes were indistinguishable in terms of WAIC (*SI Appendix*, Table S5), and we present the results for the species-varying intercept and slope model, due to slightly better model diagnostics (*SI Appendix*, Fig. S19-S22). The global relationship is given by the model: *T*_*opt*_ = −0.074 − 0.31 × *m*, where *m* is the natural log of the rescaled body mass, calculated as the species-specific ratio of mass to maturation mass.

## Discussion

In this study, we systematically analyzed the intraspecific scaling of consumption, metabolism and growth with body mass and temperature. We found strong evidence for declining optimum growth temperatures as individuals grow in size, based on two independent approaches. First, we find differences in the intraspecific mass-scaling of consumption and metabolism, and a unimodal temperature dependence of consumption, which lead to predicted declines in optimum temperature for net energy gain (and hence body growth) with size. Second, we confirm this prediction using intraspecific growth rate. Our analysis thus demonstrates the importance of understanding intraspecific scaling relationships when predicting responses of fish populations to climate warming.

That warming increases growth and development rates but reduces maximum or adult size is well known from experimental studies and is referred to as the temperature-size rule (TSR). Yet, the mechanisms underlying the TSR remain poorly understood. Pütter-type growth models, including the von Bertalanffy growth equation (VBGE), predict that the asymptotic size declines with warming if the ratio of the coefficients for energy gains and losses (*H*/*K* in Eq. 7) (Pauly & Cheung, 2018a) declines with temperature. However, the assumptions underlying the VBGE were recently questioned because of the lack of empirical basis for the scaling exponents and the effects of those on the predicted effects of temperature on asymptotic size (Lefevre et al., 2018; Marshall & White, 2019). Specifically, the allometric exponent of energy gains (*a*) is assumed to be smaller than that of energetic costs (*b*) (Eq. 7). This is based on the assumption that anabolism scales with the same power as surfaces to volumes (*a* = 2/3) and catabolism, or maintenance metabolism, is proportional to body mass (*b* = 1) (Pauly & Cheung, 2018b; von Bertalanffy, 1957). In contrast, maintenance costs are commonly thought to instead be proportional to standard metabolic rate, which in turn often is proportional to intake rates at the interspecific level (Brown et al., 2004; Marshall & White, 2019). This leads to *a* ≈ *b*, resulting in unrealistic growth trajectories and temperature dependencies of growth dynamics in Pütter models (Lefevre et al., 2018; Marshall & White, 2019). However, similar to how the existence of large fishes in tropical waters does not invalidate the hypothesis that old individuals of large-bodied fish may reach smaller sizes with warming, interspecific scaling parameters cannot reject or support these model predictions on growth *within* species. We show that the average intraspecific whole-organism mass scaling exponent of metabolism is larger than that of maximum consumption, i.e., the inequality *a* < *b* holds at the intraspecific level. By contrast, Pawar *et al*. (2012) estimated larger mass exponents for consumption than metabolic rate (0.84 and 1.04 in 2D and 3D foraging) from interspecific data, which reveals the importance of parameterizing processes occurring over ontogeny with intraspecific rather than interspecific data. When accounting for the smaller intraspecific mass exponent of consumption, and the unimodal thermal response of consumption, the thermal response of net energy gain is characterized by the optimum temperature being a function of body size (Morita et al., 2010). Therefore, empirically derived intraspecific parameterizations of simple growth models result in predictions in line with the TSR, in this case via declines in optimum growth temperatures over ontogeny rather than declines in asymptotic sizes.

Declines in optimum growth temperatures over ontogeny as a mechanism for TSR-like growth dynamics do not rely on the assumption that the ratio of the coefficients for energy gains and losses declines with temperature. In fact, we find that when using data from sub-peak temperatures only, the predicted average intraspecific activation energy of metabolism and consumption do not differ substantially, which implies there is no clear loss or gain of energetic efficiency with warming within species below temperature optima. This is in contrast to other studies, e.g. Lemoine & Burkepile (2012) and Rall *et al*. (2010). However, it is in line with the finding that growth rates increase with temperature (e.g. Angilletta & Dunham, 2003), which is difficult to reconcile from a bioenergetics perspective if warming always reduced net energy gain. Our analysis instead suggests that the mismatch between gains and losses occurs when accounting for unimodal consumption rates over temperature. The match, or mismatch, between the temperature dependence of feeding vs. metabolic rates is a central question in ecology that extends from experiments to meta-analyses to food web models (Fussmann et al., 2014; Lemoine & Burkepile, 2012; Lindmark et al., 2019; Rall et al., 2010; Vasseur & McCann, 2005). Our study highlights the importance of accounting for non-linear thermal responses for two main reasons. First, the thermal response of net energy gain reaches a peak at temperatures below the peak for consumption. Secondly, as initial warming commonly leads to increased growth rates, the effect of warming on growth rates itself depends on temperature, and growth should therefore not be assumed to be monotonically related to temperature.

Life-stage dependent optimum growth temperatures have previously been suggested as a component of the TSR (Ohlberger, 2013). Although previous studies have found declines in optimum growth temperatures with body size in some species of fishes and other aquatic ectotherms (Björnsson et al., 2007; Handeland et al., 2008; Panov & McQueen, 1998; Steinarsson & Imsland, 2003; Wyban et al., 1995), others have not (Brett et al., 1969; Elliott & Hurley, 1995). Using systematically collated growth data from experiments with variation in both size and temperature treatments (13 species), we find that for an average fish, the optimum growth temperature declines as it grows in size. This finding emerges despite the small range of body sizes used in the experiments (only 10% of observations are larger than 50% of maturation size) (*SI Appendix,* Fig. S2). Individuals of such small relative size likely invest little energy in reproduction, which suggests that physiological constraints at warmer temperatures contribute to reduced growth performance of large compared to small fish, in addition to increasing investment into reproduction (Barneche et al., 2018).

Translating results from experimental data to natural systems is challenging because maximal feeding rates, unlimited food supply, lack of predation, and constant temperatures do not reflect natural conditions, yet they affect growth rates (Brett et al., 1969; Huey & Kingsolver, 2019; Lorenzen, 1996). In addition, total metabolic costs in the wild also include additional costs for foraging and predator avoidance. It is, however, typically found that standard metabolic rate is proportional to routine metabolic rate (exhibit the same mass-scaling relationships (Kitchell et al., 1977; Messmer et al., 2017)). This assumption is also commonly applied to the relationship between maximum consumption rates and consumption rates in the wild (i.e., they are related via a constant), which has some empirical support (Neuenfeldt et al., 2020). However, the assumption of proportionality has not been tested thoroughly, despite it being identified as an important area for research over 40 years ago (Kitchell et al., 1977). If, for example, large individuals in wild populations feed more efficiently than small ones (e.g., due to being less sensitive to predation), the prediction about declining optimum growth temperature with size based on lab experiments may be altered.

Moreover, intraspecific growth rates may not appear to be unimodally related to temperature when measured over a temperature gradient across populations within a species (Denderen et al., 2020), because each population can be adapted to local climate conditions and thus display different temperature optima. However, each population likely has a thermal optimum for growth, which differs between individuals of different size. Hence, each population might have a unimodal relationship with temperature that differs from other populations of the same species. This highlights the importance of understanding the time scale of environmental change in relation to that of immediate physiological responses, acclimation, adaptation and community reorganization for the specific prediction about climate change impacts. In natural systems, climate warming may also result in stronger food limitation (Huey & Kingsolver, 2019; Ohlberger et al., 2011). Hence, as optimum growth temperatures decline not only with size but also food availability (Brett, 1971; Brett et al., 1969), and realized consumption rates often are a fraction of the maximum consumption rate (20-70%) (Kitchell *et al*. 1977; Neuenfeldt *et al*. 2020), species may be negatively impacted by warming even when controlled experiments show they can maintain growth capacity at these temperatures. Supporting this point is the observation that warming already has negative or lack of positive effects on body growth in populations living at the edge of their physiological tolerance in terms of growth (Huss et al., 2019; Neuheimer et al., 2011).

Whether the largest fish of a population will be the first to experience negative effects of warming, as suggested by the finding that optimum growth temperature declines with body size, depends on the environmental temperatures they typically experience compared to smaller conspecifics. For instance, large fish may inhabit colder temperatures compared to small fish due to ontogenetic habitat shifts (Lloret-Lloret et al., 2020; Werner & Hall, 1988); see also Heincke’s law (Audzijonyte & Pecl, 2018; Heincke, 1913). Yet, there is already empirical evidence of the largest individuals in natural populations being the first to suffer from negative impacts of warming, either from increased mortality (Peralta-Maraver & Rezende, 2021; Pörtner & Knust, 2007), or not being able to increase growth rates as smaller conspecifics tend to do (Huss et al., 2019; van Dorst et al., 2019). Hence, assuming that warming affects all individuals of a population equally is a simplification that can bias predictions of the biological impacts of climate change.

The interspecific scaling of fundamental ecological processes with body mass and temperature has been used to predict the effects of warming on body size, size structure, and population and community dynamics (Cheung et al., 2013; Gilbert et al., 2014; Morita et al., 2010; Vasseur & McCann, 2005). We argue that a contributing factor to the discrepancy between mechanistic growth models, general scaling theory, and empirical data has been the lack of data synthesis at the intraspecific level. The approach presented here can help overcome limitations of small data sets by borrowing information across species in a single modelling framework, while accounting for the intraspecific scaling of rates. Accounting for the faster increase in whole-organism metabolism than consumption with body size, the unimodal thermal response of consumption, and resulting size-dependence of optimum growth temperatures is essential for understanding what causes observed growth responses to global warming. Acknowledging these mechanisms is also important for improving predictions on the consequences of warming effects on fish growth for food web functioning, fisheries yields and global food production in warmer climates.

## Acknowledgements

We thank Hiroki Yamanaka, Dennis Tomalá Solano, Vanessa Messmer, Björn Björnsson, Albert Imsland, Tomas Árnasson, Yiping Luo, Takeshi Tomiyama and Myron Peck for generously providing data; Magnus Huss and Ken Haste Andersen, for providing useful comments on earlier versions of the manuscript; Daniel Padfield and Wilco Verberk for helpful discussions; Matthew Low and Malin Aronsson for an introduction to Bayesian inference. This study was supported by grants from the Swedish Research Council FORMAS (no. 217-2013-1315) and the Swedish Research Council (no. 2015-03752) (both to AG).

## Author contributions

ML conceived the study; ML, JO, AG designed research; ML performed research with input from JO and AG; ML, JO, AG wrote the paper and contributed to revisions of the manuscript.

## Data accessibility statement

All data and R code (lists of studies in literature search, data preparation, analyses, and figures) have been deposited in GitHub (https://github.com/maxlindmark/scaling) and Zenodo (https://doi.org/10.5281/zenodo.5806330).

