## Supporting Information for "Optimum growth temperature declines with body size within fish species"

**Appendix S1**

**Supporting Information for**

### Optimum growth temperature declines with body size within fish species

Max Lindmark[a,1], Jan Ohlberger[b], Anna Gårdmark[c]

[a] Swedish University of Agricultural Sciences, Department of Aquatic Resources, Institute of Coastal Research, Skolgatan 6, Öregrund 742 42, Sweden

[b] School of Aquatic and Fishery Sciences (SAFS), University of Washington, Box 355020, Seattle, WA 98195-5020, USA

[c] Swedish University of Agricultural Sciences, Department of Aquatic Resources, Skolgatan 6, SE-742 42 Öregrund, Sweden

[1]Author to whom correspondence should be addressed. Current address: Max Lindmark, Swedish University of Agricultural Sciences, Department of Aquatic Resources, Institute of Marine Research, Turistgatan 5, Lysekil 453 30, Sweden, Tel.: +46(0)104784137,

### Literature search, selection process and criteria

This section is an overview of the literature search approach, and below we present the search terms for each rate separately (maximum consumption, metabolism and growth). In addition to search terms, we also applied filters by selecting only the following subjects: 'marine freshwater biology', 'fisheries', 'ecology', 'zoology', 'biology', 'physiology'. For growth rates, we also included 'limnology' and for maximum consumption we included 'limnology' and 'evolutionary biology'. The use of additional subjects for growth and consumption reflects the lower data availability compared to metabolism. As we suspected that relatively few studies would have considered both body size- and temperature treatments, our goal was to get an as extensive as possible list of studies. Therefore, we also evaluated articles cited by articles found in the search, from published review-type articles and reviews of applications of bioenergetics models such as the Wisconsin model (Deslauriers et al., 2017), and if the study was found in the literature search for another rate. The source of the article (WoS search or cited in literature) is indicated in the data sets (Table S1).

Articles were filtered out at three levels of the search: title, abstract and full article. The online repository of this project (https://github.com/maxlindmark/scaling) contains .txt files of the complete list of articles found in the literature search. We removed studies from the lists if the titles made it clear the articles did not fulfil all of the following conditions: (1) experimental study, (2) fish as study organism in post-larval life stages, and (3) replicates across both body size and temperature. We treat data as individual-level rates (per fish); however, in some cases they were measured as averages across multiple individuals. In addition to these general criteria, we also had criteria specific for each rate (see below). When several studies were found for the same species, we did not include all but instead chose the study with the largest body size and temperature range (in that order), as there can be large differences in absolute values of some physiological parameters between studies.

For consumption and growth rate, we determined if each data point within species was below or beyond peak temperature for a size group within a species either by using information provided by the authors (e.g., by deriving a polynomial regression of the rate as a function of temperature to find the temperature of peak rate), by fitting Sharpe Schoolfield equations (see main text, Eq. 4) or visually inspecting data for each species separately. Whether a data point is below or above peak or optimum temperatures is indicated by a separate column in the data (Table S1).

***Maximum consumption rate***

We used the following topic terms for maximum consumption rate (three searches in total): (consumption OR bioenerg* OR ingestion OR "food-intake") AND (mass OR weight OR size) AND (temperature*), as well as: (feeding-rate OR bio-energ*) AND (mass OR weight OR size) AND (temperature*) and lastly: ("food intake") AND (mass OR weight OR size) AND (temperature*). * represents any group of characters, including no character. The searches for maximum consumption rate data resulted in 15259 articles (search date: 18 December 2018), with 3449 remaining after filtering by subject categories. The second search (search date: 13 March 2019) resulted in 431 additional titles after filtering by subject categories (of which some where duplicated from the first search) and the third search (search date: 29 June 2020) yielded 626 but no additional articles as they had either been selected already or did not meet the criteria. Articles were filtered out at the abstract and whole article stage if the original reference could not be identified and evaluated, if data were normalized (i.e., using a priori defined scaling relationships to show corrected data rather than measured values), there was no acclimation, or if measurements were not maximum consumption rate. As with the growth data, definitions of ad-libitum feeding may differ between studies – the key for our purpose is that food rations led to satiation and were not limiting. Consumption rates were converted to $g\ day^{-1}$. These data were compiled in the file consumption_data.xlsx.

***Metabolic rate***

We used the following topic terms for metabolic rate data: (metabolism OR "oxygen-consumption" OR "oxygen consumption") AND (mass OR weight OR size) AND (temperature*). * represents any group of characters, including no character. The search for metabolic rate experiments resulted in 8405 articles (search date: 6 June 2019), which was reduced to 3458 after applying filters for subject categories. Articles were filtered out at the abstract and whole article stage if the original reference could not be identified and evaluated, if data were normalized (i.e., using a priori defined scaling relationships to normalize data for data a given size rather than measured values), if there was no acclimation or if it was not standard, routine or resting metabolic rate. The latter was defined as oxygen consumption of an unfed fish at no or little spontaneous activity. Metabolic rates were converted to $mg\ O_2\ h^{-1}$, because it was the most common unit in the data set. These data were compiled in the file metabolism_data.xlsx.

***Growth rates & optimum temperature for growth over body mass***

Growth rates were taken from data found in the literature search for optimum growth temperatures. Therefore, articles in which growth rates were measured at sub-optimum temperatures only were not included (note this is in contrast to consumption data where “optimum” was not included in the search terms). We used the following topic terms for growth rate data: (growth) AND (mass OR weight OR size) AND (temperature*) AND (optimum), as well as: (growth) AND (mass OR weight OR size) AND (temperature*) AND (optim*). * represents any group of characters, including no character. The two searches for growth rates resulted in 3313 articles (search date: 22 March 2019), and 3747 articles (search date: 5 May 2019), respectively. After applying additional filters by subject category, we acquired 566 and 893 studies, respectively (of which some are duplicates due to similar search-strings). We removed studies at the abstract and whole article stage where the original reference could not be identified and evaluated, if we could not extract actual growth rates, if there was not a controlled temperature for each growth trial, or if there were not multiple defined size-classes. We used only one observation (data point) per size class and temperature treatment, and in cases where there were two, we used the mean value. In addition, we ensured that no other treatment (e.g., food limitation) confounded the response variable and thus only used data from experiments with satiating food levels. Body mass is either the geometric mean of the initial and final mass of the growth trial or the size class (as reported by the original authors), depending on data availability (see Table S1). It is important to control for feeding rations as it affects the temperature optimum for growth (Brett et al., 1969). This was achieved in different ways in the different experimental studies, but commonly involved excess feeding rations once or several times per day. The key description we looked for in the study was that food was not limiting. We treat data as individual-level growth (per fish); however, these were commonly measured as averages for multiple individuals. In the case growth was length-based, we converted it to mass using weight-length relationships from FishBase (Froese et al., 2014; Froese & Pauly, 2019). We compiled two separate data sets: raw growth rates (growth_data.xlsx) and temperature at optimum growth (growth_data_Topt.xlsx). In the latter, we defined optimum temperature for growth as the fitted optimum temperature by size-class (usually estimated in the original study). Therefore, the optimum temperature may not always correspond to an actual experimental temperature but could be an estimation between two temperature treatments. If the optimum temperature (by size group) was not estimated in the original study, we used the temperature where growth rate was maximized. All growth rates were expressed in unit % day$^{-1}$.

**Table S1** Explanation of data columns (G=growth data, $T_{opt}$=optimum growth temperature data, C=maximum consumption data, M=metabolism data).

| Column | Explanation | Datasets |
|---|---|---|
| ***growth_rate_%/day*** | Main response variable. | G, $T_{opt}$ |
| ***opt_temp_c*** | Main response variable. | $T_{opt}$ |
| ***initial_mass_g*** | Body mass [g] at the onset of the growth trial. | G, $T_{opt}$ |
| ***final_mass_g*** | Body mass [g] at the end of the growth trial. | G, $T_{opt}$ |
| ***geom_mean_mass_g*** | Geometric mean mass in $t_1$ and $t_2$ of the growth trial. | G, $T_{opt}$ |
| ***size_group*** | Representative body mass of size group in the growth trial, in case initial, final or geometric body mass could not be retrieved. | G, $T_{opt}$ |
| ***consumption*** | Main response variable. | C |
| ***metabolic_rate*** | Main response variable. | M |
| ***type*** | Type of respiration measurement (resting, routine, standard). | M |
| ***unit*** | Unit of response variable. | C, M |
| ***original_unit*** | Original unit of response variable. If different from "***unit***", see "***notes***" column for information on conversion. | C, M |
| ***mass_g*** | Body mass in experiment [g]. Some studies report body masses before and some after the feeding trials. See "***notes***". | C, M |
| ***temp_c*** | Experimental temperature [℃]. | G, C, M |
| ***above_peak_temp*** | Is the experiment conduced at temperature above peak temperature for the given size group? Y/N. | G, C, M |
| ***common_name*** | Common name of species. | G, $T_{opt,}$ C, M |
| ***species*** | Scientific name of species. | G, $T_{opt,}$ C, M |
| ***genus*** | Genus of species. | G, $T_{opt,}$ C, M |
| ***family*** | Family of species. | G, $T_{opt,}$ C, M |
| ***order*** | Order of species. | G, $T_{opt,}$ C, M |
| ***habitat*** | Species natural habitat, taken from FishBase | G, $T_{opt,}$ C, M |

| | | |
|---|---|---|
| | (Froese & Pauly, 2019). | |
| ***lifestyle*** | Lifestyle of species, taken from FishBase (Froese & Pauly, 2019). | G, $T_{opt}$, C, M |
| ***biogeography*** | Biogeography of species, taken from FishBase (Froese & Pauly, 2019). | G, $T_{opt}$, C, M |
| ***trophic_level*** | Trophic level of species, taken from FishBase (Froese & Pauly, 2019). | G, $T_{opt}$, C, M |
| ***w_maturation_g*** | Body mass [g] at maturation taken from FishBase (Froese & Pauly, 2019). If not available, weight was estimated from length using species-specific allometric weight-length, else taken from alternative sources (see ”***notes***”). Used to estimate relative body size across species in the data and to normalized optimum growth temperatures across species. | G, $T_{opt}$ |
| ***w_max_published_g*** | Max. published weight [g] taken from FishBase (Froese & Pauly, 2019). If not available, weight was estimated from length using species-specific allometric weight-length, else taken from alternative sources (see ”***notes***”). Used to estimate relative body size across species in the data. | G, $T_{opt}$, C, M |
| ***env_temp_min*** | Min. environmental temperature [°C], taken from FishBase (Froese & Pauly, 2019). If not available on FishBase, data were taken from alternative sources (see ”***notes***”). Used to compare experimental temperatures to common temperatures for species. | G, $T_{opt}$, C, M |
| ***env_temp_max*** | Max. environmental temperature [°C], taken from FishBase (Froese & Pauly, 2019). If not available on FishBase, data were taken from alternative sources (see ”***notes***”). Used to compare experimental temperatures to common temperatures for species. | G, $T_{opt}$, C, M |
| ***env_temp_mid*** | Median of environmental temperature [°C], taken from FishBase (Froese & Pauly, 2019). If not available on FishBase, data were taken from alternative sources (see ”***notes***”). Used to compare experimental temperatures to common temperatures for species. | G, $T_{opt}$, C, M |
| ***pref_temp_mid*** | Median of preferred temperature [°C], taken from FishBase (Froese & Pauly, 2019). If not available on FishBase, data were taken from alternative sources (see ”***notes***”). Used to compare experimental temperatures to common temperatures for species. | G, $T_{opt}$, C, M |
| ***notes*** | This column contains additional information, including if data were sent by authors, if any column above has data that is not from the main source (i.e. FishBase), how certain metrics were calculated, alternative common names, comments | G, $T_{opt}$, C, M |

| | | |
|---|---|---|
| | on the experimental protocol, information on conversion to standard "***unit***", source of the data (literature search or cited in paper from literature search) | |
| ***reference*** | Source (See Table S2). | G, $T_{opt,}$ C, M |

**Table S2** Species, common name, the data set(s) in which they appear and the sources (G=growth data, $T_{opt}$=optimum growth temperature data, C=maximum consumption data, M=metabolism data). If more than one data and source, the sources are in order (1 study per species and rate).

| Species | Common name | Datasets | Source |
|---|---|---|---|
| ***Pseudopleuronectes yokohamae*** | Marbled flounder | G, $T_{opt}$, C | (Tomiyama et al., 2018) |
| ***Cyclopterus lumpus*** | Lumpfish | G, $T_{opt}$ | (Nytrø et al., 2014) |
| ***Paralichthys olivaceus*** | Japanese flounder (alt. bastard halibut, Japaneese halibut or Olive flounder) | G, $T_{opt}$, C | (Iwata et al., 1994) |
| ***Salvelinus alpinus*** | Arctic char | G, $T_{opt}$ | (Siikavuopio et al., 2013) |
| ***Salmo salar*** | Atlantic salmon | G, $T_{opt}$ | (Handeland et al., 2008) |
| ***Lates calcarifer*** | Barramundi | G, $T_{opt}$, C, M | (Bermudes et al., 2010)<br>(Bermudes et al., 2010)<br>(Bermudes et al., 2010)<br>(Glencross & Felsing, 2006) |
| ***Gadus morhua*** | Atlantic cod | G, $T_{opt}$, M | (Björnsson et al., 2007)(Tirsgaard et al., 2015) |
| ***Hippoglossus hippoglossus*** | Atlantic halibut | G, $T_{opt}$ | (Björnsson & Tryggvadóttir, 1996) |
| ***Scophthalmus maximus*** | Turbot | G, $T_{opt}$ | (Árnason et al., 2009) |
| ***Boreogadus saida*** | Arctic cod | G, $T_{opt}$ | (Laurel et al., 2017) |
| ***Rachycentron canadum*** | Cobia | G, $T_{opt}$, C | (Sun & Chen, 2014) |
| ***Pelteobagrus fulvidraco*** | Yellow catfish | G, $T_{opt}$,C | (Zhang et al., 2017) |
| ***Anarhichas minor*** | Spotted wolffish | G, $T_{opt}$ | (Imsland et al., 2006) |
| ***Oncorhynchus mykiss*** | Rainbow trout | C, M | (From & Rasmussen, 1984) |
| ***Perca fluviatilis*** | Eurasian perch | C | (Lessmark, 1983) |

| | | | |
|---|---|---|---|
| ***Phoxinus phoxinus*** | Eurasian minnow | C, M | (Cui & Wootton, 1988) |
| ***Coregonus hoyi*** | Bloater | C | (Binkowski & Rudstam, 1994) |
| ***Pomoxis annularis*** | White crappie | C | (Hayward & Arnold, 1996) |
| ***Gambusia affinis*** | Western mosquitofish | C | (Chipps & Wahl, 2004) |
| ***Morone saxatilis*** | Striped bass | C | (Duston et al., 2004) |
| ***Salvelinus fontinalis*** | Brook trout | C, M | (Baldwin, 1957) (Beamish, 1964) |
| ***Leuciscus leuciscus*** | Dace | C | (Marmulla & Rosch, 1990) |
| ***Lepomis microlophus*** | Redear sunfish | C | (Wang et al., 2003) |
| ***Channa argus*** | Chinese snakehead (alt. Northern snakehead or Snakehead) | C, M | (Liu et al., 1998) (Liu et al., 2000) |
| ***Siniperca chuatsi*** | Mandarin fish | C, M | (Liu et al., 1998) (Liu et al., 2000) |
| ***Gasterosteus aculeatus*** | Three-spined stickleback | C | (Wootton et al., 1980) |
| ***Salmo trutta*** | Brown trout | C | (Elliott, 1976) |
| ***Epinephelus coioides*** | Orange-spotted grouper | C | (Lin et al., 2008) |
| ***Coregonus albula*** | Vendace | M | (Ohlberger et al., 2012) |
| ***Coregonus fontanae*** | Stechlin cisco | M | (Ohlberger et al., 2012) |
| ***Abramis brama*** | Common bream | M | (Ohlberger et al., 2012) |
| ***Rutilus rutilus*** | Common roach | M | (Ohlberger et al., 2012) |
| ***Salvelinus confluentus*** | Bull trout | M | (Mesa et al., 2013) |
| ***Catostomus commersonii*** | White sucker | M | (Beamish, 1964) |
| ***Cyprinus carpio*** | Common carp | M | (Beamish, 1964) |
| ***Ameiurus nebulosus*** | Brown bullhead | M | (Beamish, 1964) |
| ***Silurus meridionalis*** | Southern catfish | M | (Xie & Sun, 1990) |
| ***Carassius auratus*** | Goldfish | M | (Beamish & Mookherjii, 1964) |
| ***Pomadasys commersonnii*** | Spotted grunter | M | (Du Perez et al., 1986) |
| ***Melanogrammus aeglefinus*** | Haddock | M | (Peck et al., 2005) |
| ***Centropristis striata*** | Black sea bass | M | (Slesinger et al., 2019) |
| ***Anguilla anguilla*** | European eel | M | (Degani et al., 1989) |
| ***Micropterus salmoides*** | Largemouth bass | M | (Glover et al., 2012) |
| ***Cyprinodon macularius*** | Desert pupfish | M | (Heuton et al., 2018) |
| ***Micropogonias undulatus*** | Atlantic croaker | M | (Horodysky et al., 2011) |
| ***Leiostomus xanthurus*** | Spot | M | (Horodysky et al., |

| | | | |
|---|---|---|---|
| | | | 2011) |
| ***Coreius guichenoti*** | Largemouth bronze gudgeon | M | (Luo & Wang, 2012) |
| ***Sprattus sprattus*** | European sprat | M | (Meskendahl et al., 2010) |
| ***Plectropomus leopardus*** | Leopard coral grouper | M | (Messmer et al., 2017) |
| ***Galaxias maculatus*** | Common galaxias | M | (Milano et al., 2016) |
| ***Polyodon spathula*** | American paddlefish (alt. Mississippi paddlefish) | M | (Patterson et al., 2013) |
| ***Argyrosomus japonicus*** | Mulloway | M | (Pirozzi & Booth, 2009) |
| ***Lythrypnus dalli*** | Bluebanded goby | M | (Rangel & Johnson, 2018) |
| ***Colossoma macropomum*** | Tambaqui (alt. Cachama) | M | (Tomala et al., 2014) |
| ***Carassius auratus grandoculis*** | Round crucian carp (alt. Nigorobuna) | M | (Yamanaka et al., 2013) |

### Data overview

***Maximum consumption & metabolic rate***

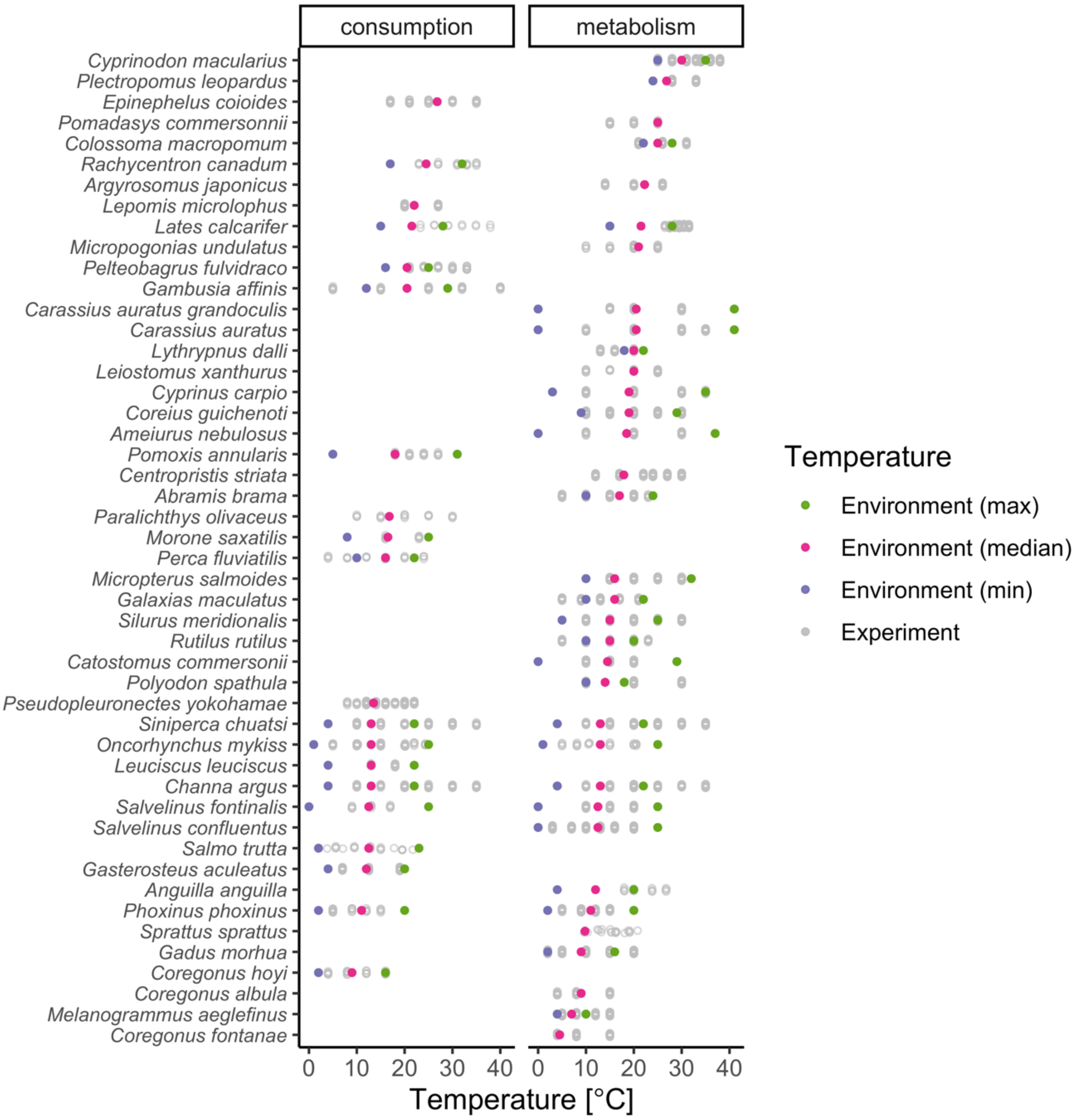

*Fig. S1. Experimental temperatures (gray) and environmental (min, median and max) temperatures (purple, pink and green, respectively) of species represented in the consumption (left) and metabolism (right) data sets. Missing temperatures means information was not available on FishBase. Experimental temperatures are jittered vertically for visibility.*

***Growth rate***

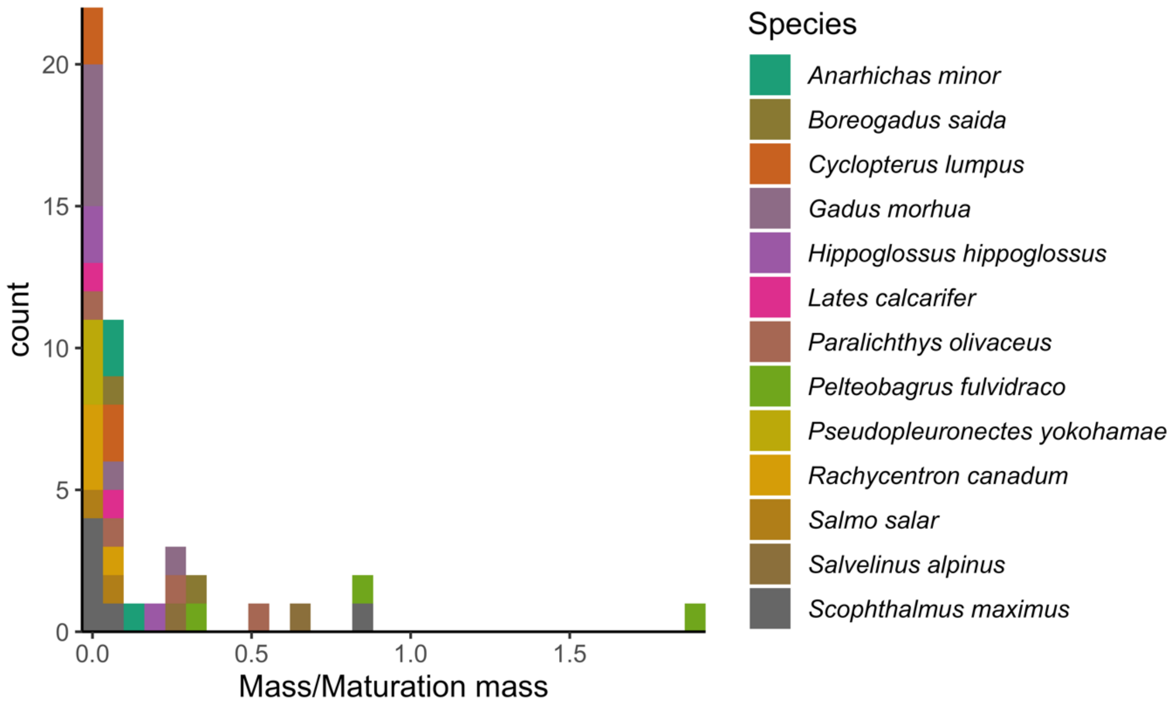

*Fig. S2. The distribution of rescaled masses for individual observations (mass/mass at maturation), where color indicate species.*

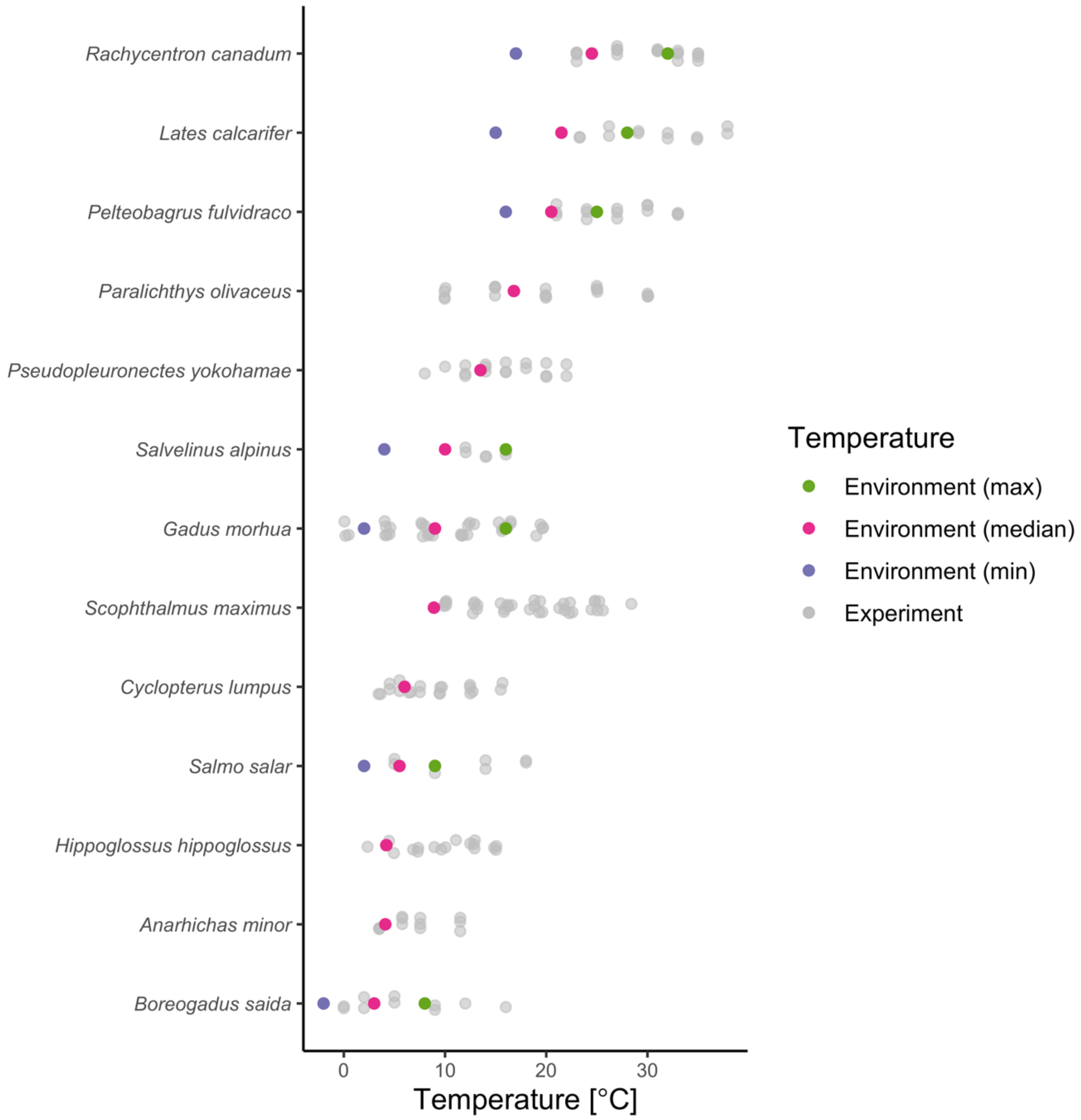

*Fig. S3. Experimental temperatures (gray) in the growth rate data and environmental (min, median and max) temperatures (purple, pink and green, respectively). Missing temperatures means information was not available on FishBase. Experimental temperatures are jittered vertically for visibility.*

### Supplementary methods and analysis

**Table S3** Description of model parameters (type and their interpretation in brackets) and their prior distributions (see '*Model description'* and equations 1-3 in the main text). *N* refers to a normal distribution (mean and standard deviation, s.d.) and *U* to a uniform distribution (interval). For simplicity, only the parameters of the full model are shown here (i.e., with most coefficients varying by species), but note that when a model is fitted with a common rather than species-varying coefficient, for example $\beta_1$ instead of $\beta_{1j} \sim N(\mu_{\beta_1}, \sigma_{\beta_1})$, we use the same prior for $\beta_1$ as for $\mu_{\beta_1}$.

| Model | Parameter | Description | Prior distribution |
|---|---|---|---|
| **Log-linear regressions for consumption and metabolism** | $\mu_{\beta_{0s}}$ | Hyperparameter (average intercept for standard metabolic rate across species). *Only for the metabolism models.* | $N(-2, 5)$ |
| | $\mu_{\beta_{0r}}$ | Hyperparameter (average intercept for routine and resting metabolic rate across species). *Only for the metabolism models.* | $N(-1, 5)$ |
| | $\mu_{\beta_0}$ | Hyperparameter (average intercept across species). *Only for the consumption models.* | $N(0, 5)$ |
| | $\mu_{\beta_1}$ | Hyperparameter (average mass coefficient across species). | $N(0.75, 1)$ |
| | $\mu_{\beta_2}$ | Hyperparameter (average temperature coefficient across species) | $N(-0.6, 1)$ |
| | $\mu_{\beta_3}$ | Hyperparameter (average interaction coefficient across species) | $N(0, 1)$ |
| | $\sigma_{\beta_{0s}}$ | Hyperparameter (s.d. of species-intercepts for standard metabolic rate) | $U(0, 10)$ |
| | $\sigma_{\beta_{0r}}$ | Hyperparameter (s.d. of species-intercepts for routine and resting metabolic rate) | $U(0, 10)$ |
| | $\sigma_{\beta_1}$ | Hyperparameter (s.d. of species mass coefficients) | $U(0, 10)$ |
| | $\sigma_{\beta_2}$ | Hyperparameter (s.d. of species temperature coefficients) | $U(0, 10)$ |
| | $\sigma_{\beta_3}$ | Hyperparameter (s.d. of species interaction coefficients) | $U(0, 10)$ |
| | $\sigma$ | Parameter (s.d.) | $U(0, 10)$ |
| **Sharpe-Schoolfield (unimodal consumption data)** | $\mu_{C_{0j}}$ | Hyperparameter (average consumption at reference temperature [-10 on centered scale] across species) | $N(1, 1)$ |
| | $\mu_{E_j}$ | Hyperparameter (average activation energy across species) | $N(0.5, 0.5)$ |
| | $E_h$ | Parameter (common rate of decline with temperature) | $N(2, 2)$ |

| | | | |
|---|---|---|---|
| | $T_h$ | Parameter (common temperature at which half the rate is reduced due to high temperatures) | $N(5, 2)$ |
| | $\sigma_{E_j}$ | Hyperparameter (s.d. of species-varying activation energies) | $U(0, 3)$ |
| | $\sigma_{C_{0j}}$ | Hyperparameter (s.d. of species-varying average consumption) | $U(0, 3)$ |
| | $\sigma$ | Parameter (s.d.) | $U(0, 3)$ |
| **Linear $\boldsymbol{T_{opt}}$ models** | $\mu_{\beta_0}$ | Hyperparameter (average intercept across species) | $N(0, 5)$ |
| | $\mu_{\beta_1}$ | Hyperparameter (average mass coefficient across species) | $N(0, 5)$ |
| | $\sigma_{\beta_0}$ | Hyperparameter (s.d. of species-intercepts) | $U(0, 10)$ |
| | $\sigma_{\beta_1}$ | Hyperparameter (s.d. of species mass coefficients) | $U(0, 10)$ |
| | $\sigma$ | Parameter (s.d.) | $U(0, 10)$ |

**Table S4**. Model comparison for the log-linear regressions of how consumption and metabolism depend on mass and temperature below optimum temperatures (see '*Model description'* and equations 1-3 in the main text). The column m*t indicates whether the model for the rate includes an interactive effect of mass and temperature. The models differ in which coefficients vary among species and which are common, where $\beta_0$ is the intercept, $\beta_1$ mass coefficient (mass-exponent on linear scale), $\beta_2$ temperature coefficient (corresponding to the negative activation energy) and $\beta_3$ interaction between mass and temperature. The WAIC columns shows ΔWAIC and absolute WAIC in brackets, rounded to the nearest decimal, where ΔWAIC is the difference between each model's WAIC and the lowest WAIC across models. Bold indicates models with ΔWAIC < 2.

| Model | m*t | Species-varying parameter(s) | WAIC metabolism | WAIC consumption |
|---|---|---|---|---|
| **M1** | Yes | $\beta_0, \beta_1, \beta_2, \beta_3$ | **0 (274.6)** | 3.4 (564.6) |
| **M2** | | $\beta_0, \beta_1, \beta_2$ | **1.2 (275.8)** | 1.87 (563.1) |
| **M3a** | | $\beta_0, \beta_1$ | 305.8 (580.4) | 147.1 (708.3) |
| **M3b** | | $\beta_0, \beta_2$ | 385.3 (659.9) | 68.5 (629.7) |
| **M4** | | $\beta_0$ | 648.6 (923.2) | 189.4 (750.6) |
| **M5** | No | $\beta_0, \beta_1, \beta_2$ | 6.1 (280.6) | **0 (561.2)** |
| **M6a** | | $\beta_0, \beta_1$ | 348.2 (622.8) | 164.8 (726.0) |
| **M6b** | | $\beta_0, \beta_2$ | 386.6 (661.2) | 72.5 (633.7) |
| **M7** | | $\beta_0$ | 681.5 (956.1) | 211.9 (773.1) |

**Table S5**. Comparison of the two models fitted to optimum growth temperature data. The WAIC columns shows ΔWAIC and absolute WAIC in brackets, rounded to the nearest decimal, where ΔWAIC is the difference between each models' WAIC and the lowest WAIC across models. Bold indicates models with ΔWAIC < 2.

| **Model** | **Species-varying parameter(s)** | **WAIC** |
|---|---|---|
| **M1** | $\beta_0, \beta_1$ | **0 (177.3)** |
| **M2** | $\beta_0$ | **1 (178.3)** |

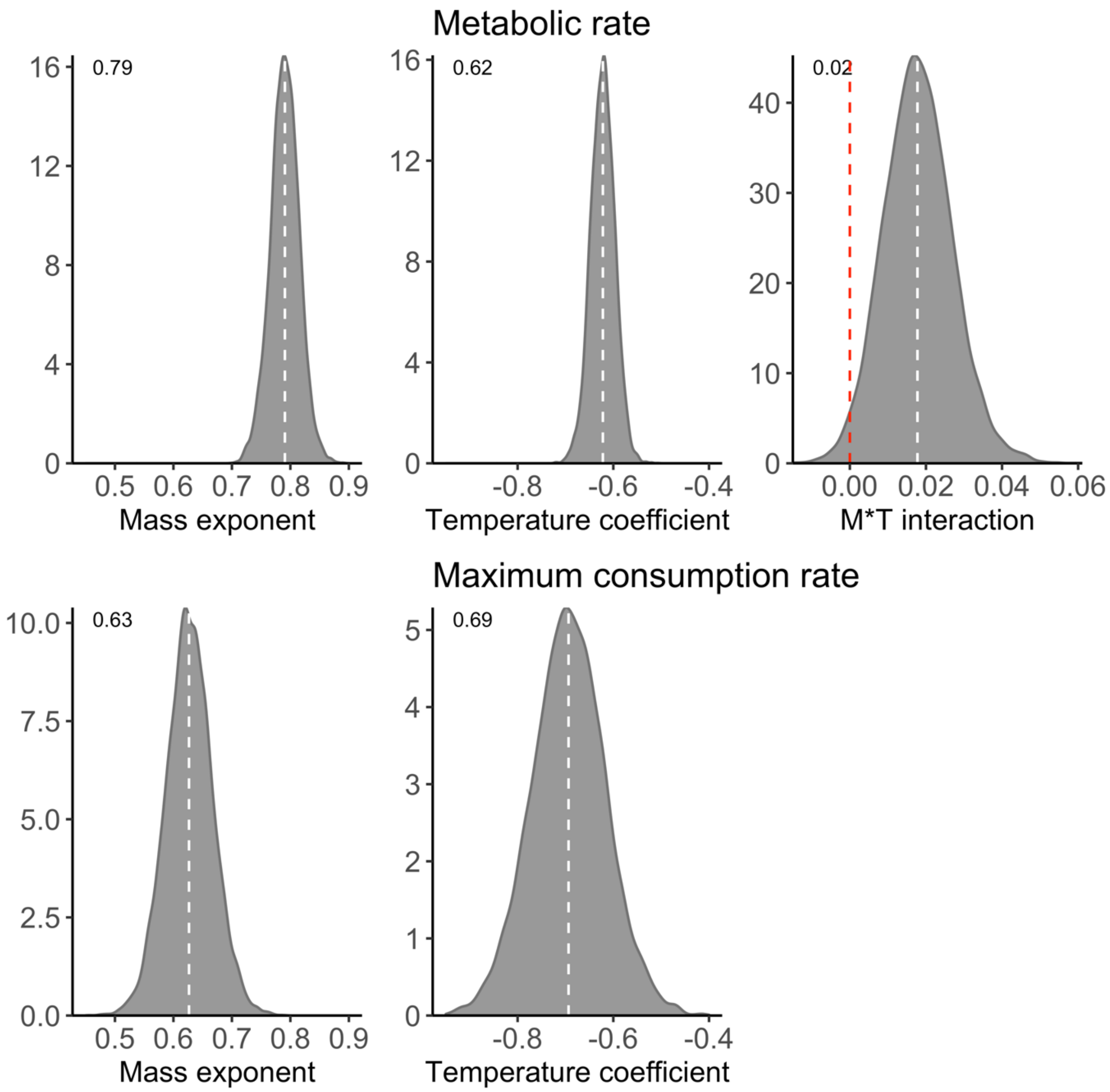

*Fig. S4. Posterior distributions of the global intraspecific mass exponents ($\mu_{\beta_1}$) and temperature coefficients ($\mu_{\beta_2}$) for metabolic rate (top) and maximum consumption rate (bottom). For metabolism, the global interaction coefficient ($\mu_{\beta_3}$) is also shown (estimated and presented on an Arrhenius temperature scale), but for consumption this term was not included in the selected model. Numbers in the top left corner correspond to the posterior median. The axes are the same for each parameter for comparison between the two rates.*

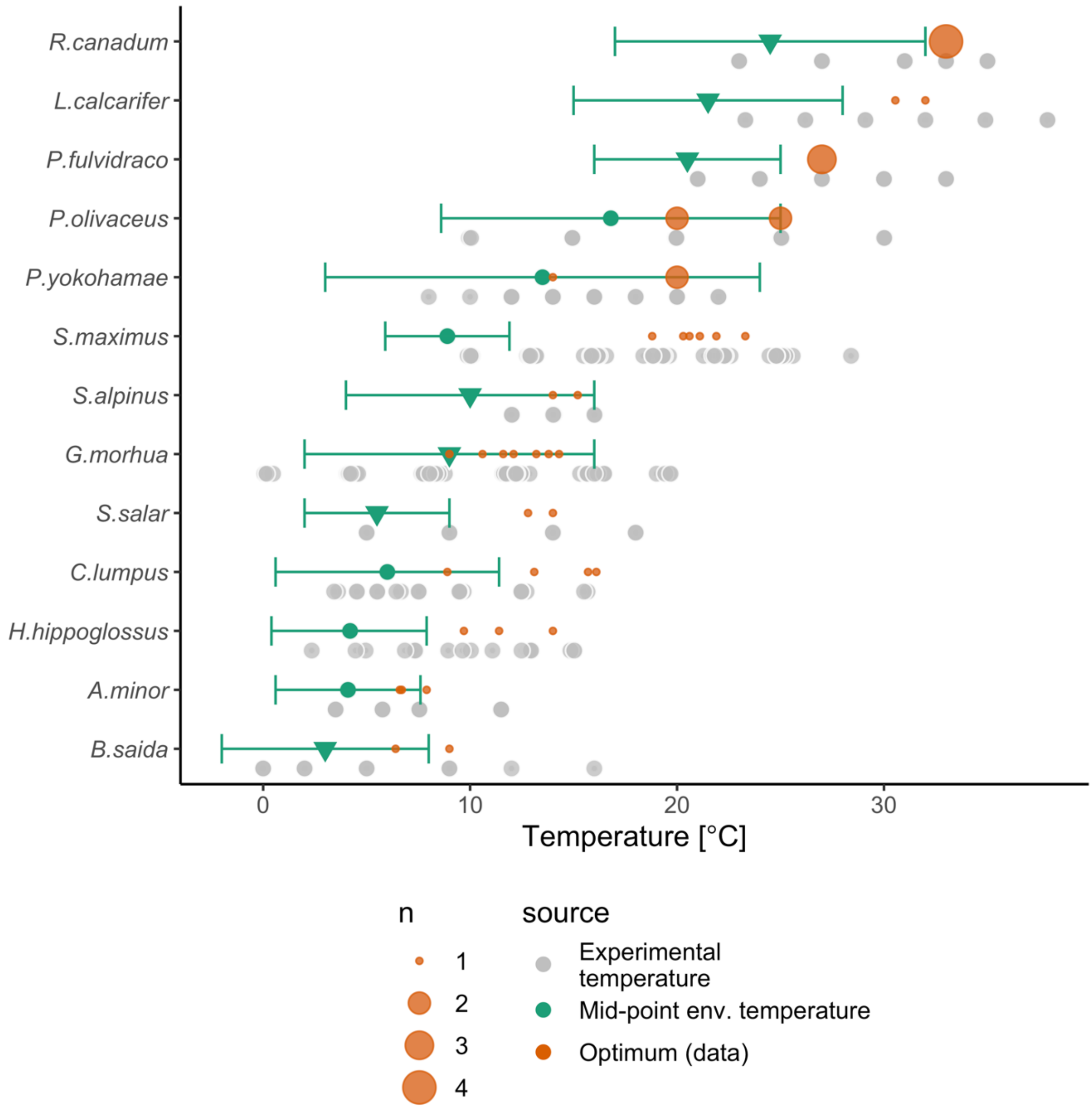

*Fig. S5. Experimental temperatures (grey) overlap environmental temperatures (green), and optimum growth temperatures (orange) are typically at the upper end or above the environmental range. Horizontal green lines show the minimum and maximum environmental temperature based on either temperature in distribution range (triangles) or modelled distribution maps (circles), both taken from FishBase. The optimum growth temperatures are depicted for all size-classes per species, where the circle size is proportional to number of observations at that temperature.*

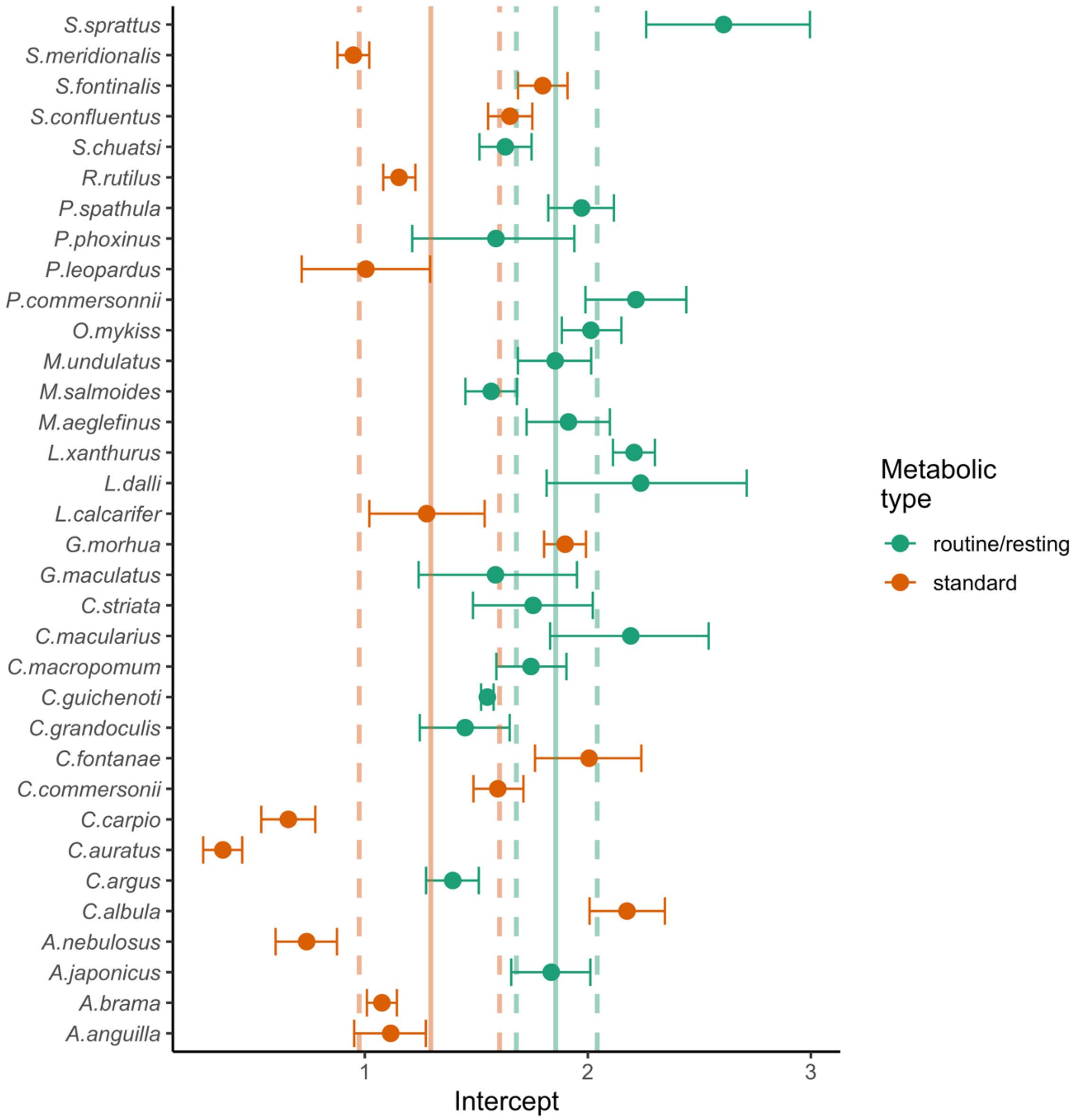

*Fig. S6. Posterior median of species-level intercepts (points) and their 95% credible interval (horizontal error bars). Colors indicate the type of metabolism measurement for each species. Vertical solid lines are the posterior medians of the global intercepts (orange for standard metabolic rate, $\mu_{\beta_{0s}}$, and green for routine or resting metabolic rate, $\mu_{\beta_{0r}}$), and the dashed vertical lines show the 95% credible intervals for the global parameters.*

#### Model validation and fit

Figures showing convergence of species-level parameters can be found on: [https://github.com/maxlindmark/scaling](https://github.com/maxlindmark/scaling), in this section only global parameters are visualized.

***Maximum consumption rate – below peak temperatures***

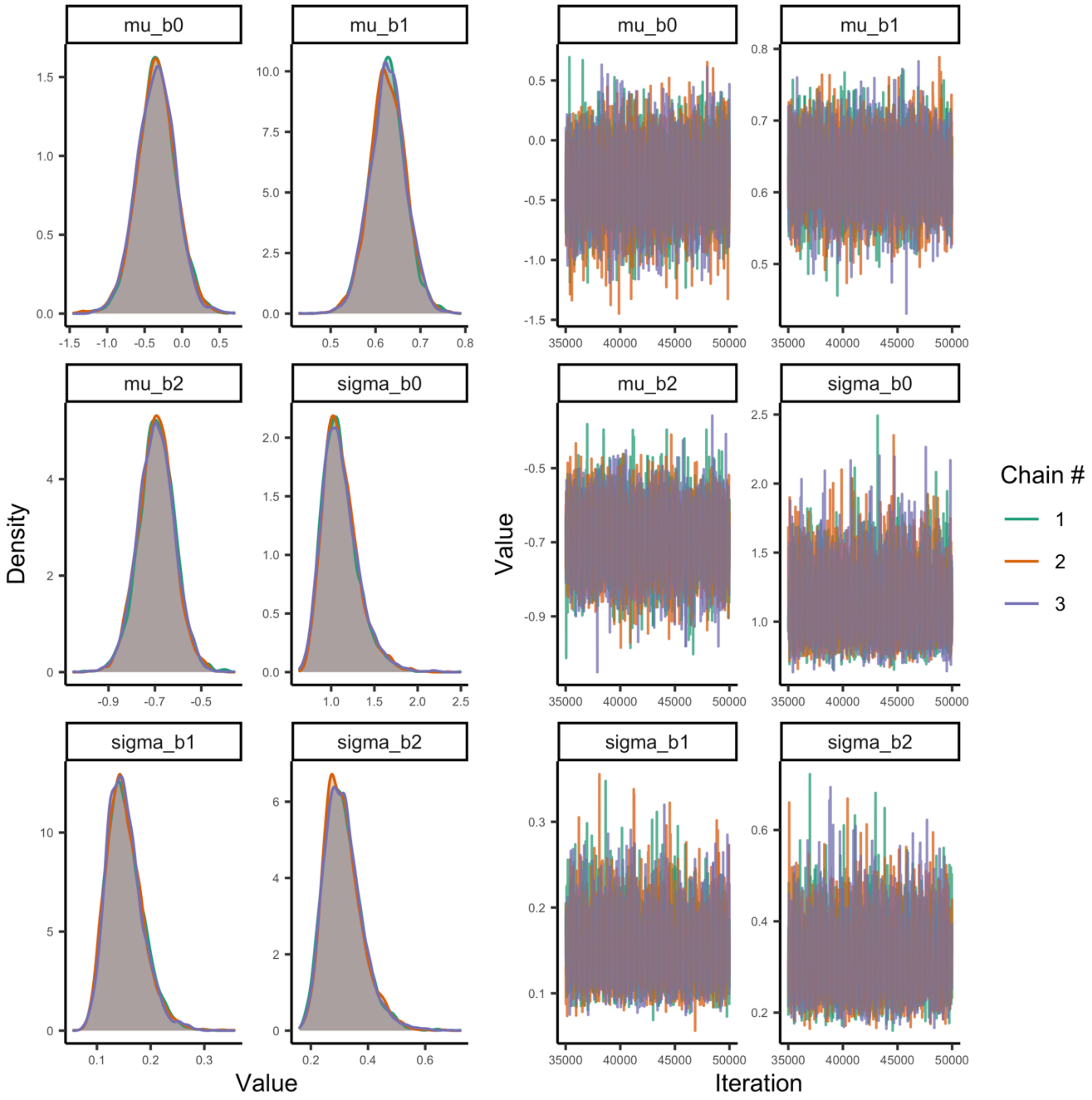

*Fig. S7. Posterior densities and trace plots for evaluation of chain convergence (by chain, indicated by color), for the global-level parameters for the log-linear maximum consumption rate model at temperatures below peak temperatures.*

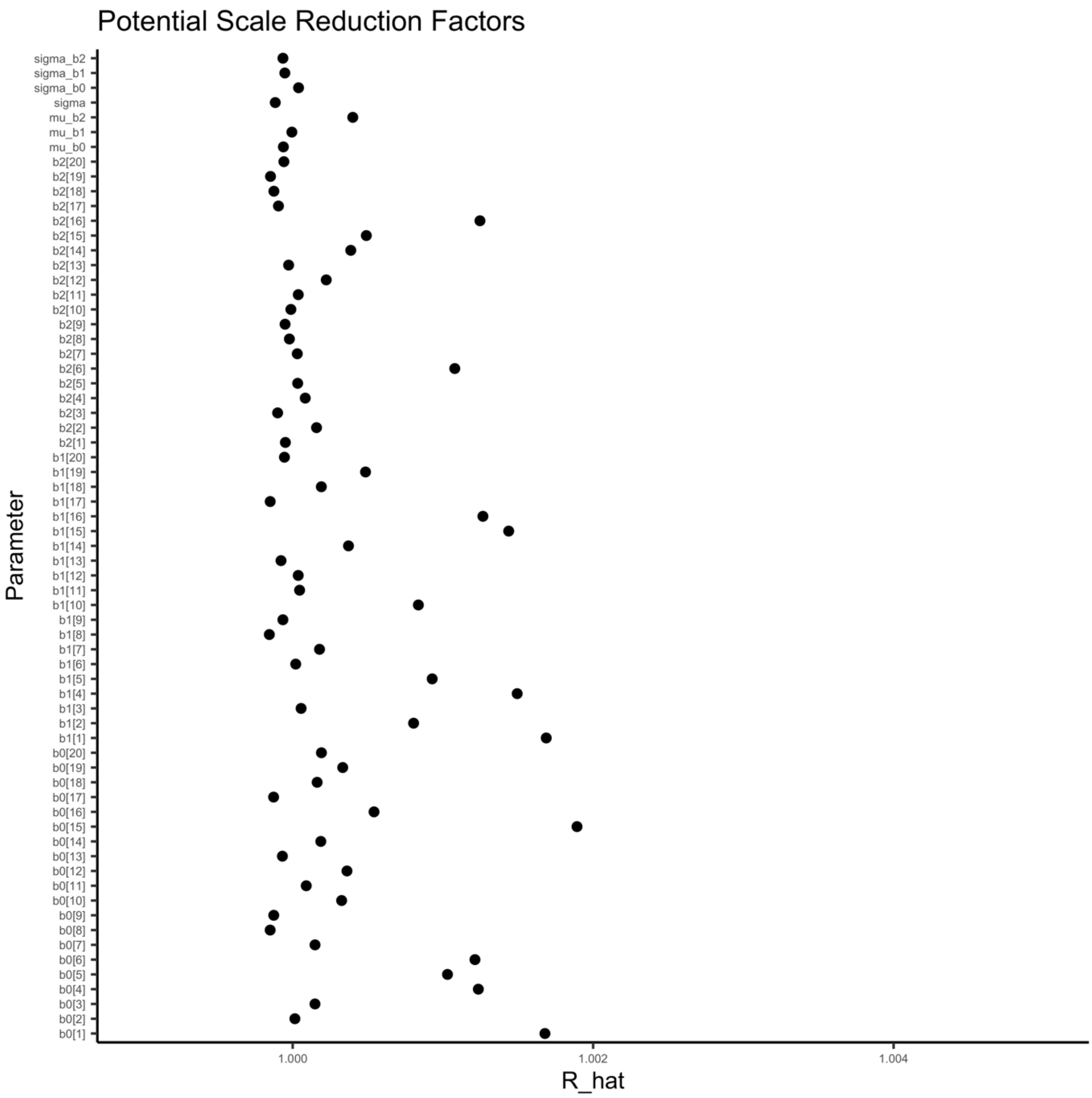

*Fig. S8. Potential scale reduction factor ($\hat{R}$) for the log-linear maximum consumption rate model. This factor is based on the comparison of between and within-chain variation for the same parameter. A value close to one implies chains converged to the same distribution. The index of the parameter corresponds to species in alphabetical order.*

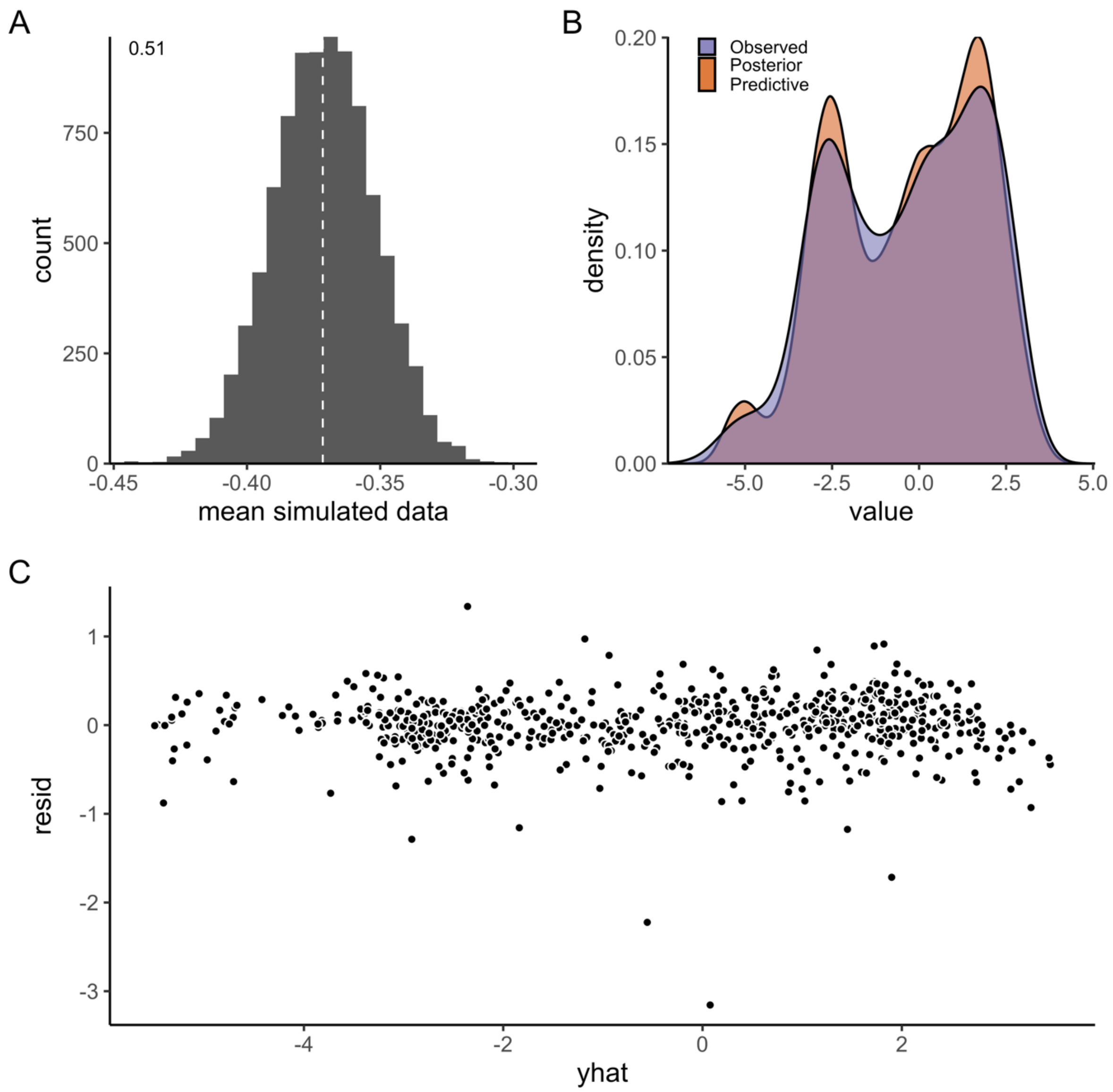

*Fig. S9. A) Model fit (mean) for the log-linear model of maximum consumption rate at temperatures below temperature peak (by species). Fit is evaluated by simulating data from the likelihood (at each iteration of the MCMC chain), to compare how well it matches the original data. Each simulated data point is assigned a 0 or 1 if it is below or above the mean data point (the vertical line corresponds to the mean in data). The number in the plot corresponds to the mean of the vector of 0's and 1's. B) Posterior predictive distribution (orange) and distribution of data (purple). C) Difference between the observed value and the posterior median of the predicted value, plotted against fitted value.*

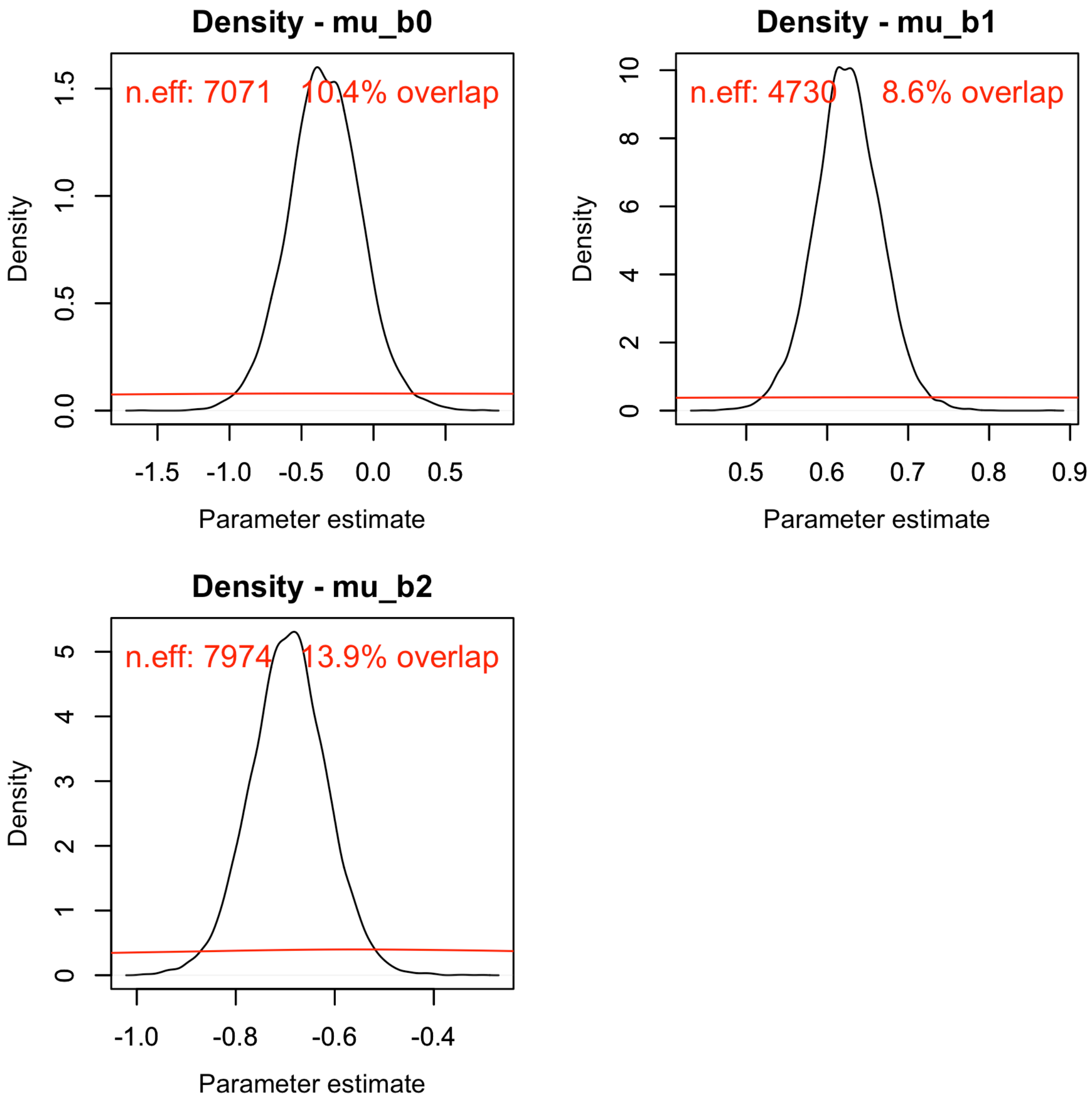

*Fig. S10. Posterior (black) and prior distribution (red) for the global parameters in the log-linear model for maximum consumption rate, including their % overlap and effective sample size (n.eff).*

***Maximum consumption rate – including beyond peak temperatures***

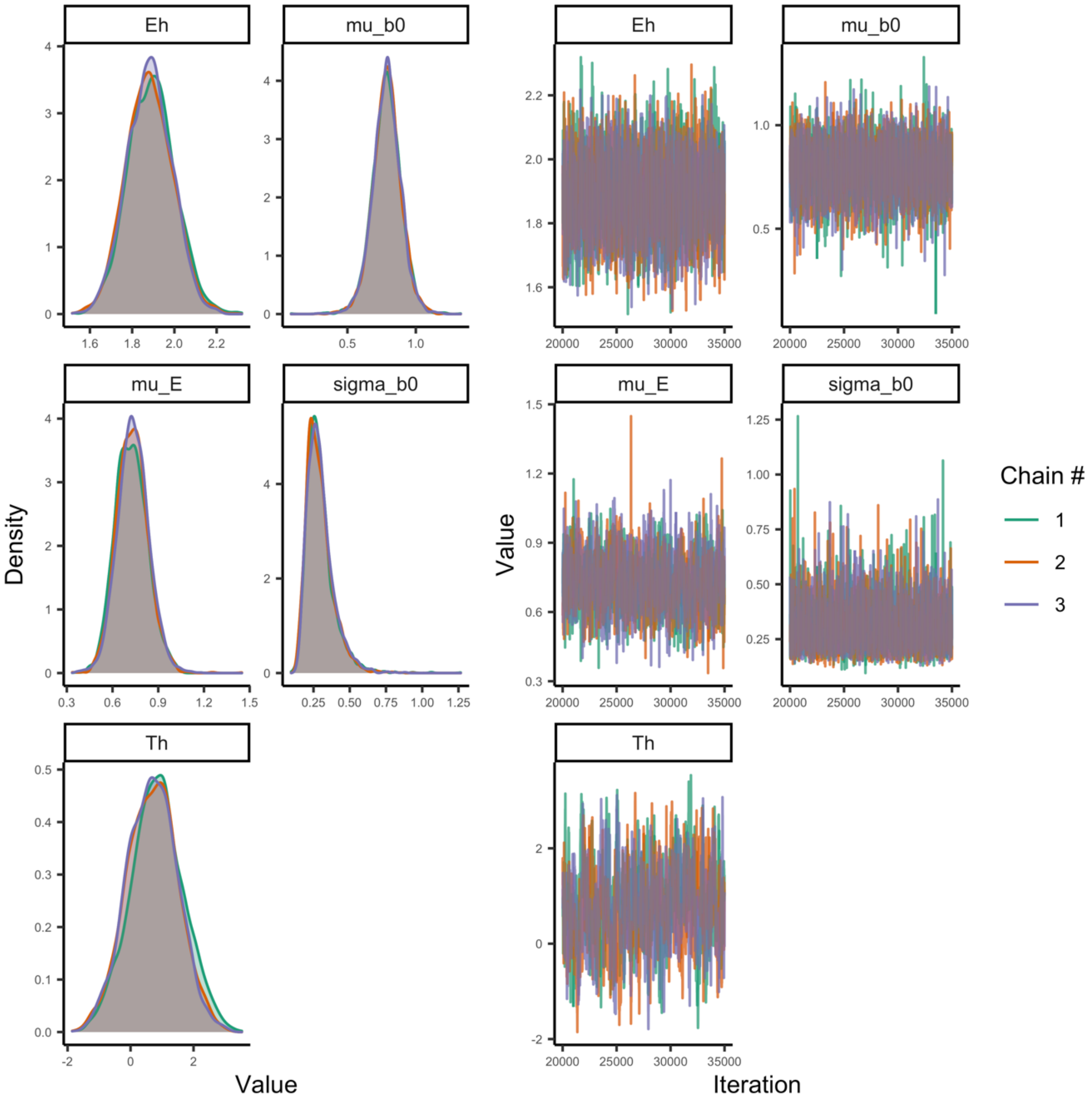

*Fig. S11. Posterior densities and trace plots for evaluation of chain convergence (by chain, indicated by color), for the global-level parameters for the Sharpe-Schoolfield model fitted to maximum consumption rate data with temperatures including beyond peak temperatures.*

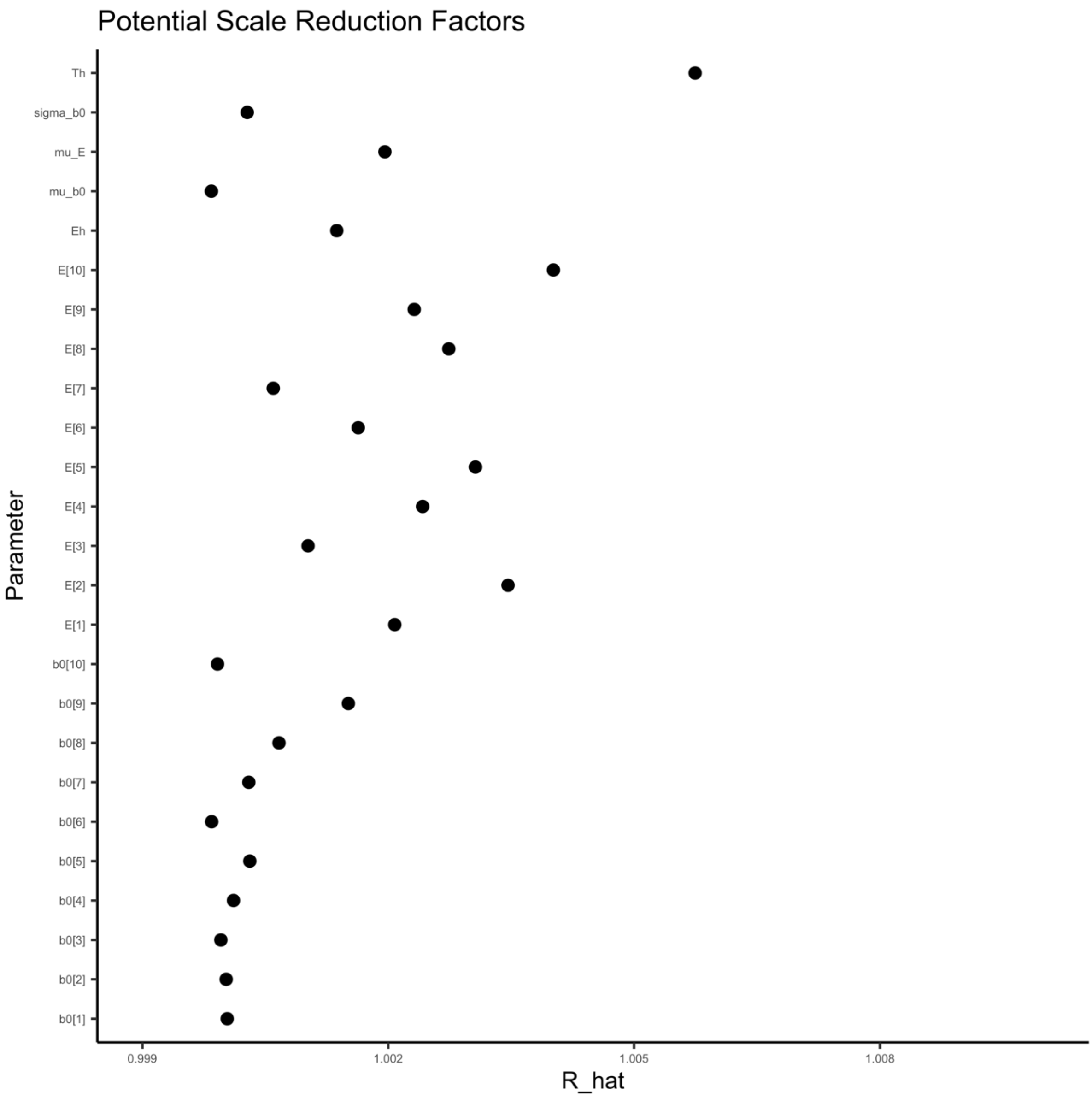

*Fig. S12. Potential scale reduction factor ($\hat{R}$) for the Sharpe-Schoolfield model fitted to maximum consumption rate data (including data beyond peak). This factor is based on the comparison of between and within-chain variation for the same parameter. A value close to one implies chains converged to the same distribution. The index of the parameter corresponds to species in alphabetical order.*

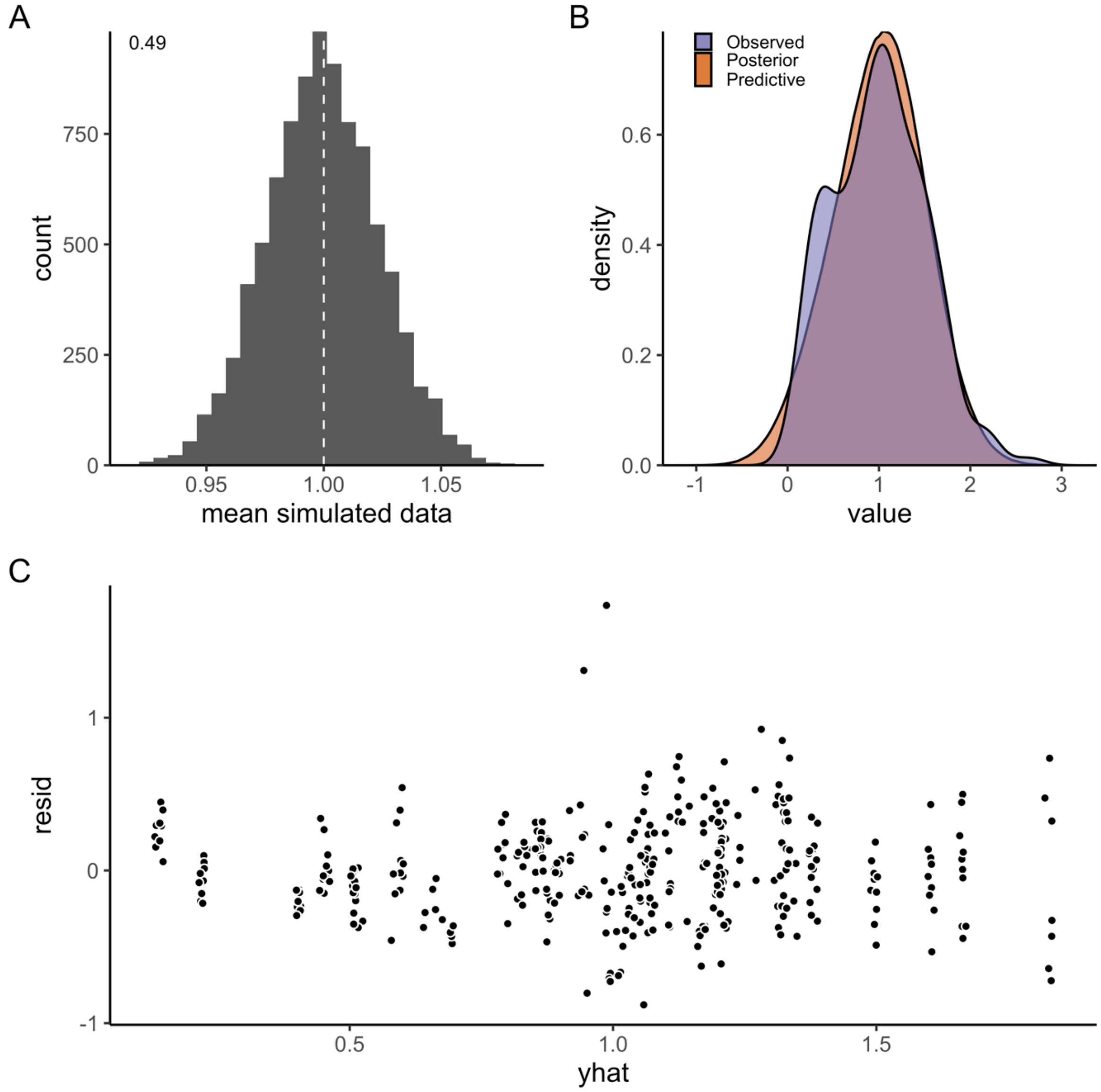

*Fig. S13. A) Model fit (mean) for the Sharpe-Schoolfield model fitted to maximum consumption rate data including temperatures beyond peak (by species). Fit is evaluated by simulating data from the likelihood (at each iteration of the MCMC chain), to compare how well it matches the original data. Each simulated data point is assigned a 0 or 1 if it is below or above the mean data point (the vertical line corresponds to the mean in data). The number in the plot corresponds to the mean of the vector of 0's and 1's. B) Posterior predictive distribution (orange) and distribution of data (purple). C) Difference between the observed value and the posterior median of the predicted value, plotted against fitted value.*

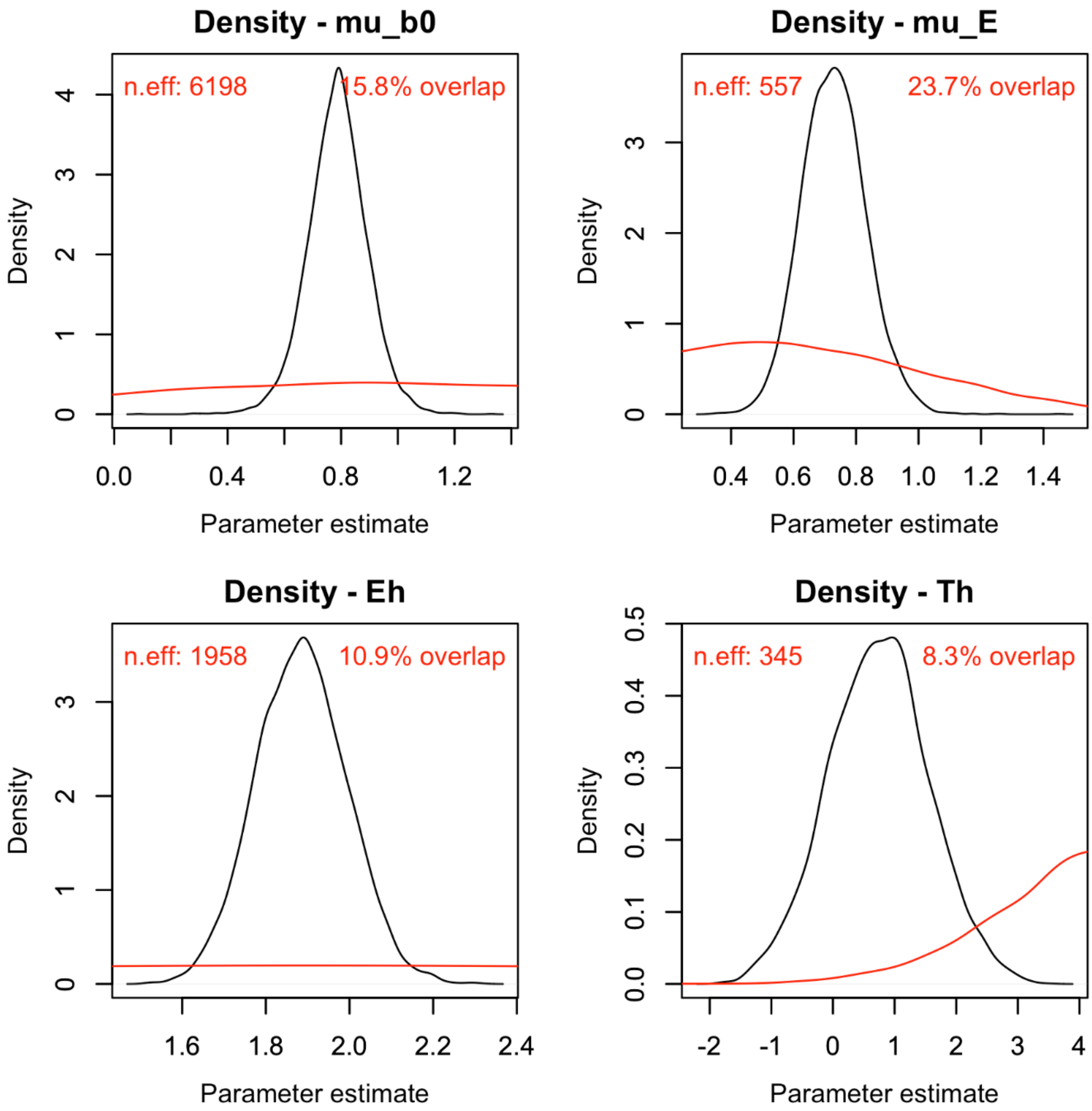

*Fig. S14. Posterior (black) and prior distribution (red) for the global parameters in the Sharpe-Schoolfield model for maximum consumption rate including data beyond peak, including their % overlap (rounded) and effective sample size (n.eff).*

***Metabolic rate***

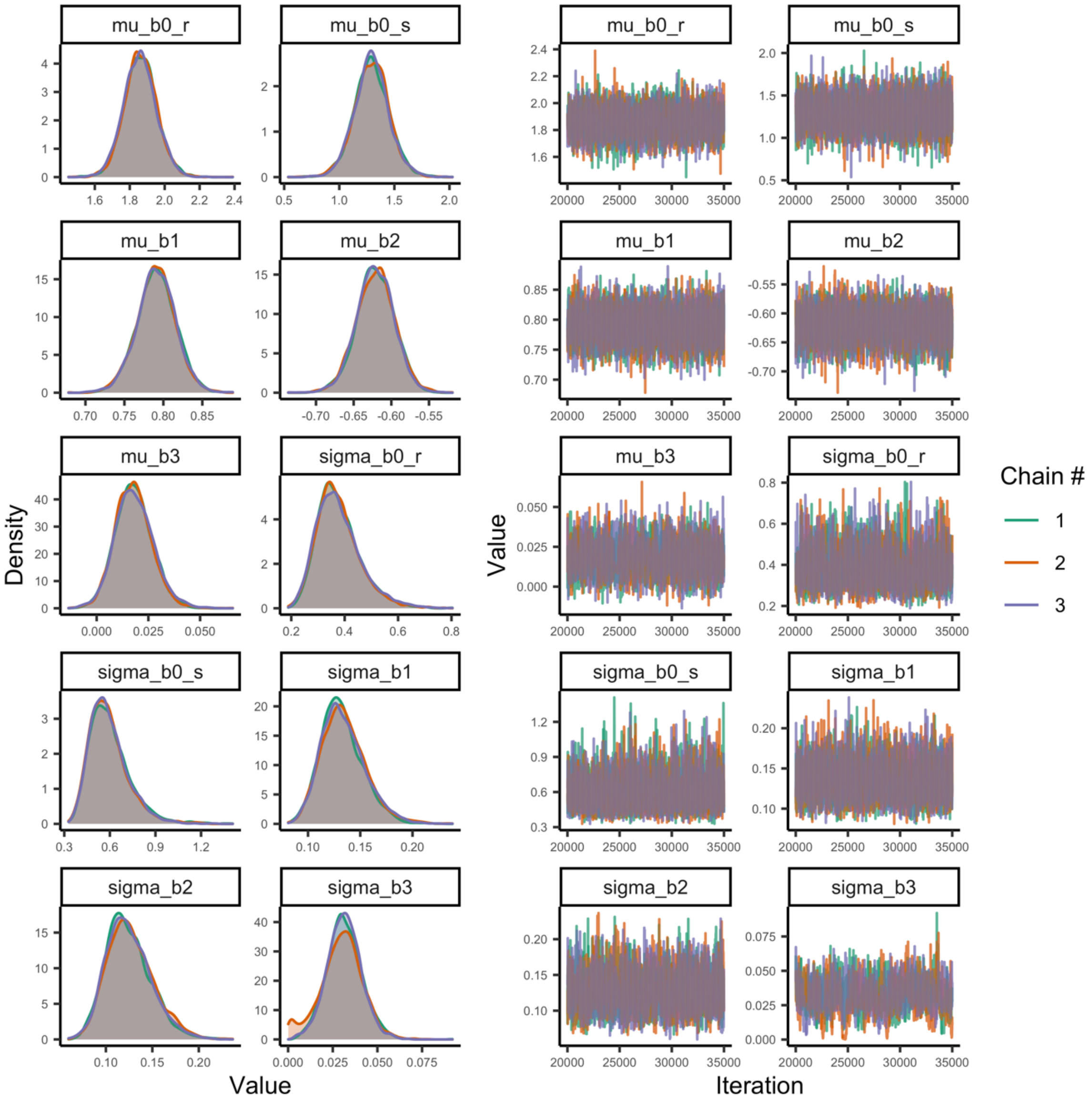

*Fig. S15. Posterior densities and trace plots for evaluation of chain convergence (by chain, indicated by color), for the global-level parameters for the metabolic rate model at temperatures below peak temperatures.*

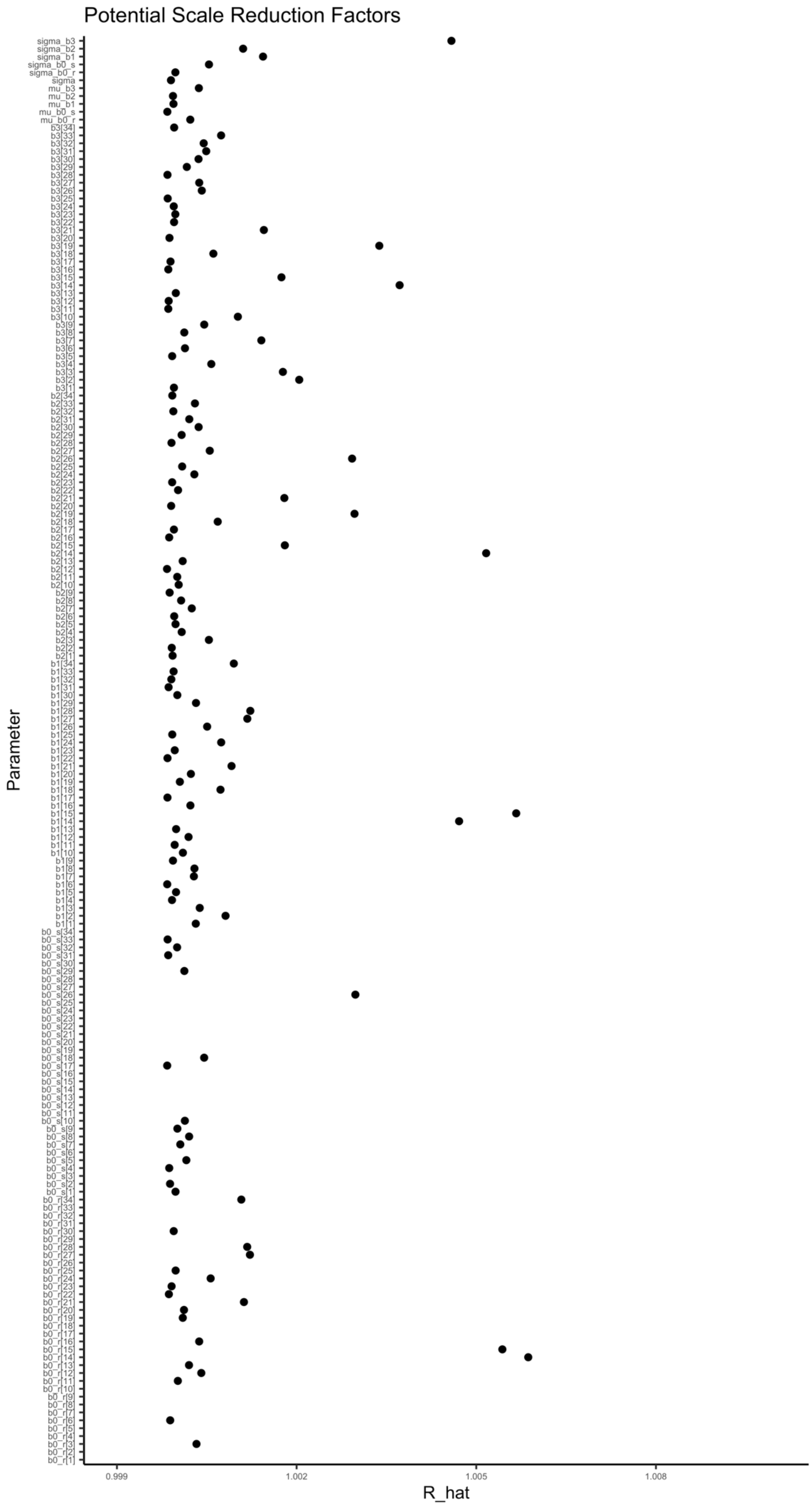
Potential Scale Reduction Factors
Parameter
R_hat
0.999
1.002
1.005
1.008

*Fig. S16. Potential scale reduction factor ($\hat{R}$) for the metabolic rate model. This factor is based on the comparison of between and within-chain variation for the same parameter. A value close to one implies chains converged to the same distribution. The index of the parameter corresponds to species in alphabetical order. Note that species with routine metabolism do not have estimates for a standard metabolic rate intercept and vice versa, hence, not all parameters in the graph have $\hat{R}$ values.*

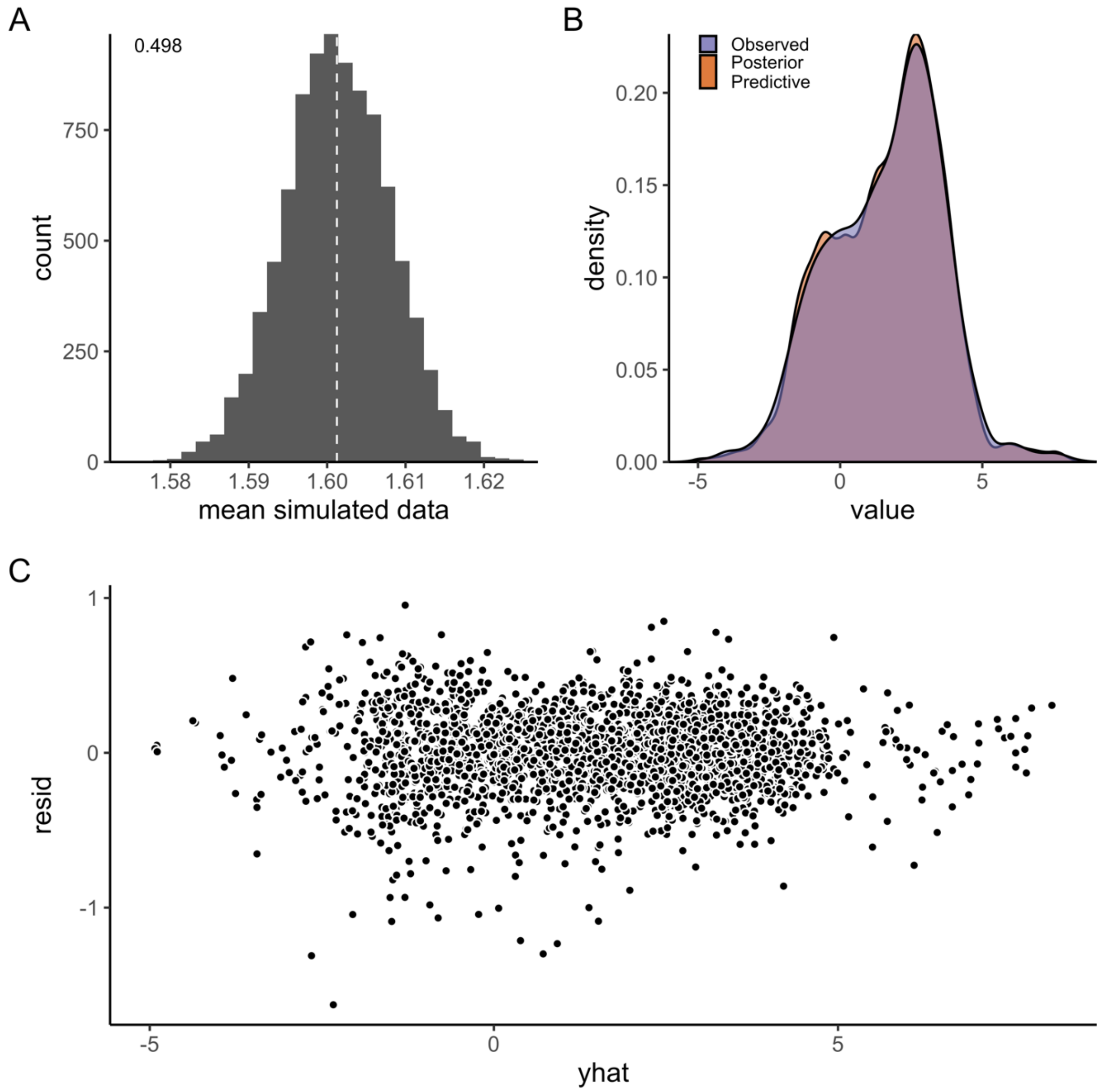

*Fig. S17. A) Model fit (mean) for the log-linear model of metabolic rate. Fit is evaluated by simulating data from the likelihood (at each iteration of the MCMC chain), to compare how well it matches the original data. Each simulated data point is assigned a 0 or 1 if it is below or above the mean data point (the vertical line corresponds to the mean in data). The number in the plot corresponds to the mean of the vector of 0's and 1's. B) Posterior predictive distribution (orange) and distribution of data (purple). C) Difference between the observed value and the posterior median of the predicted value, plotted against fitted value.*

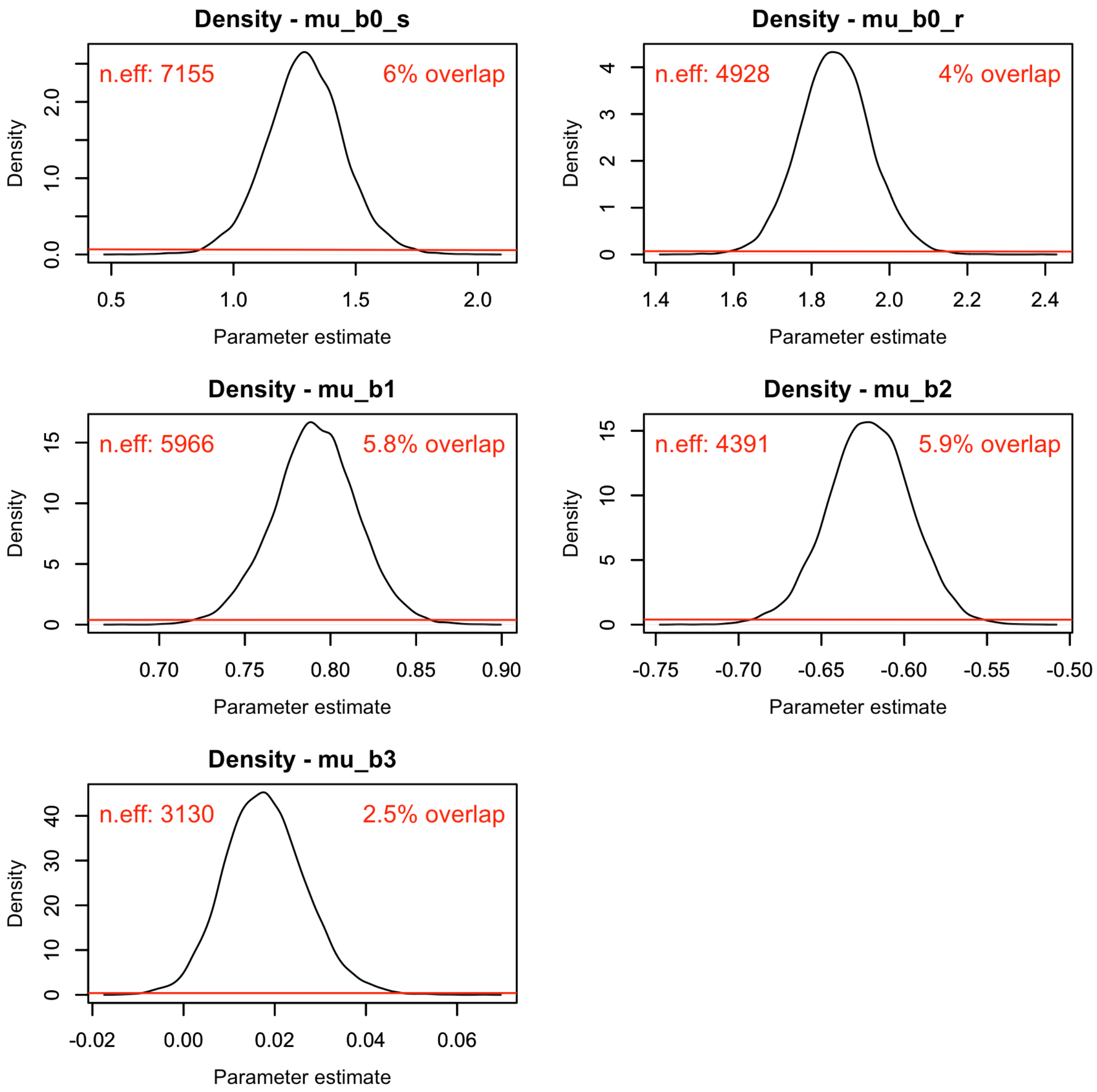

*Fig. S18. Posterior (black) and prior distribution (red) for the global parameters in the model for metabolic rate, including their % overlap and effective sample size (n.eff).*

***Optimum growth temperature***

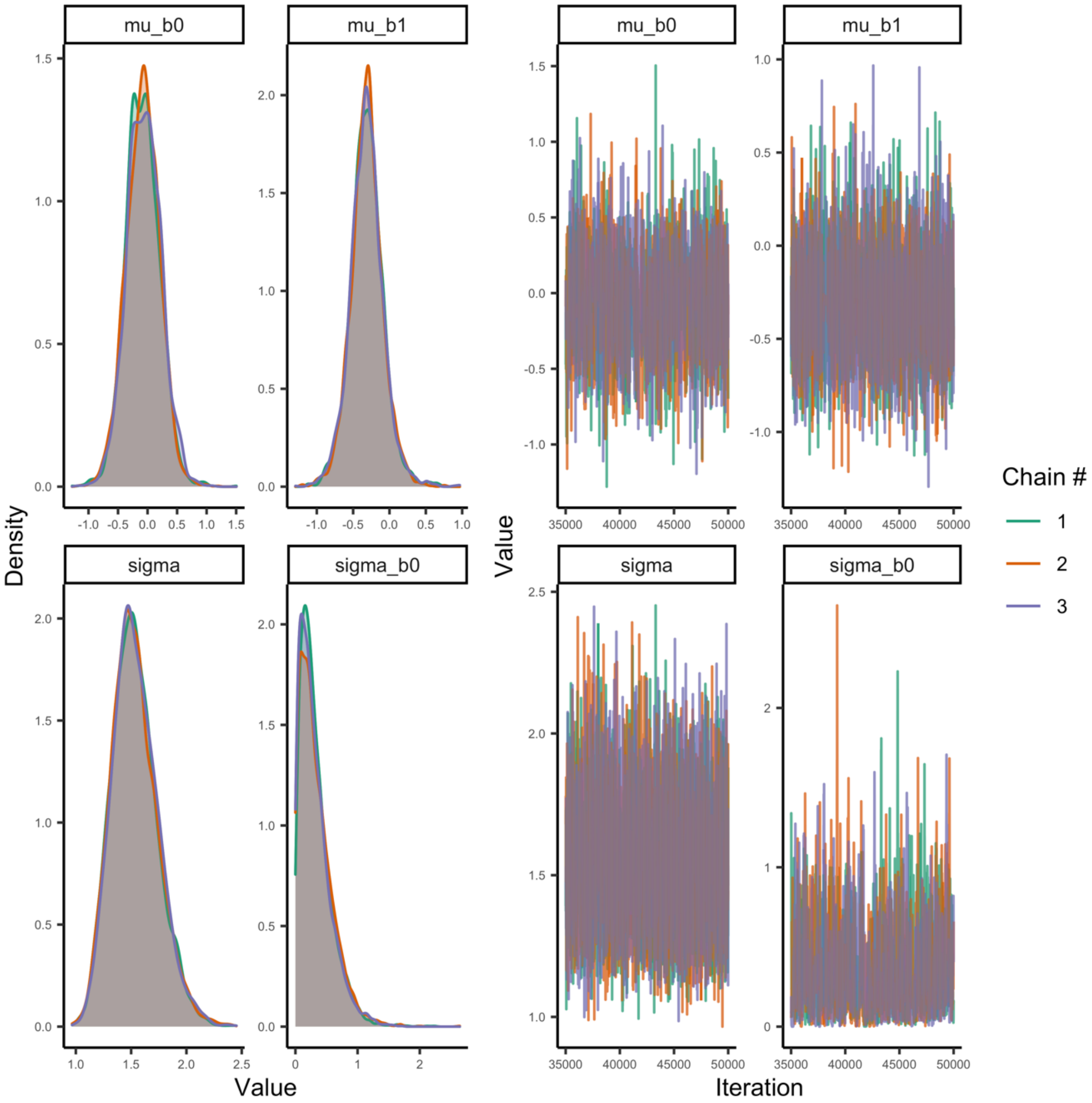

*Fig. S19. Posterior densities and trace plots for evaluation of chain convergence (by chain, indicated by color), for the global-level parameters for the* $T_{opt}$ *model.*

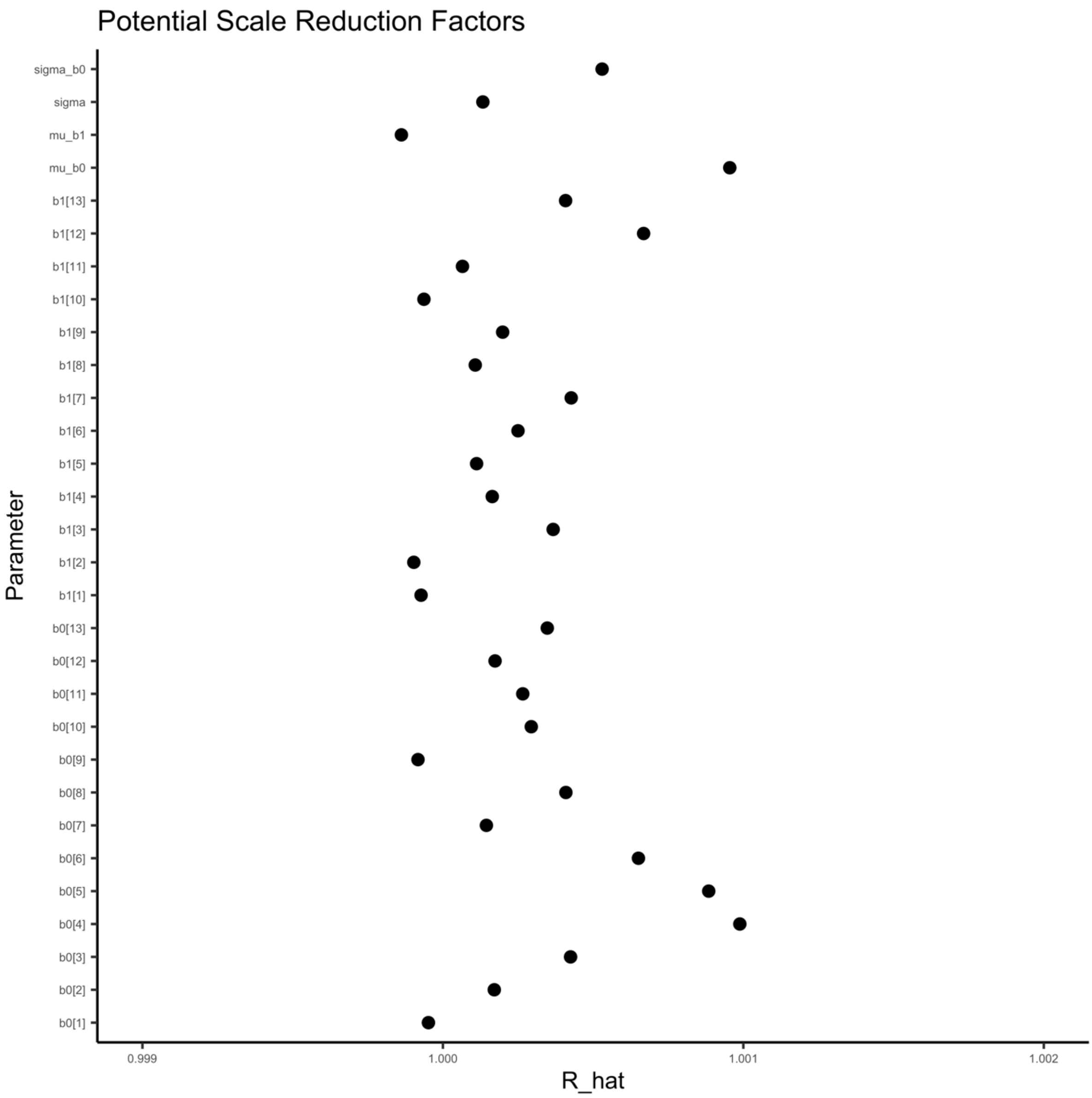

*Fig. S20. Potential scale reduction factor ($\hat{R}$) for the $T_{opt}$ model. This factor is based on the comparison of between and within-chain variation for the same parameter. A value close to one implies chains converged to the same distribution. The index of the parameter corresponds to species in alphabetical order.*

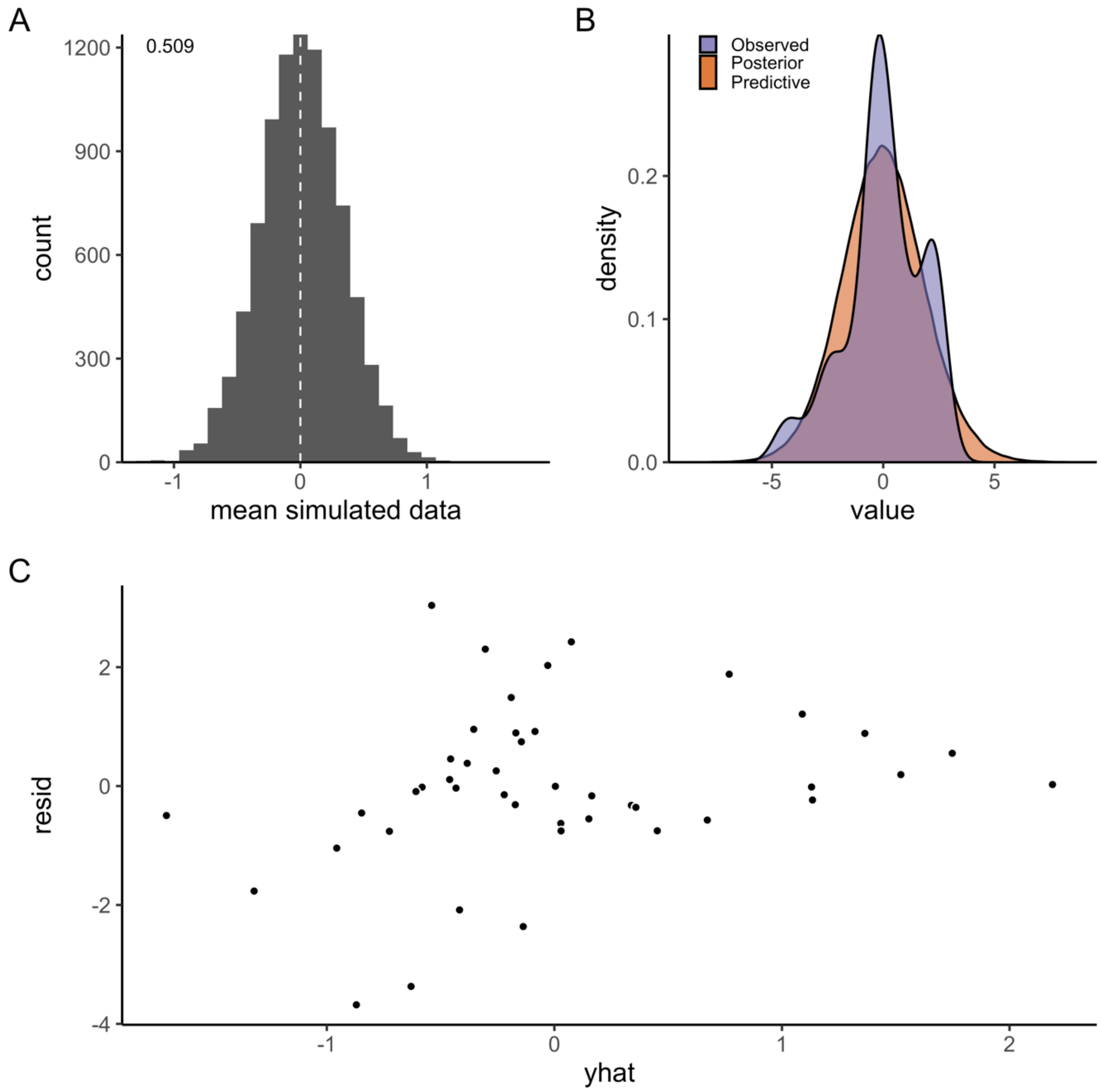

*Fig. S21. A) Model fit (mean) for the model of optimum growth temperature as a function of body mass. Fit is evaluated by simulating data from the likelihood (at each iteration of the MCMC chain), to compare how well it matches the original data. Each simulated data point is assigned a 0 or 1 if it is below or above the mean data point (the vertical line corresponds to the mean in data). The number in the plot corresponds to the mean of the vector of 0's and 1's. B) Posterior predictive distribution (orange) and distribution of data (purple). C) Difference between the observed value and the posterior median of the predicted value, plotted against fitted value.*

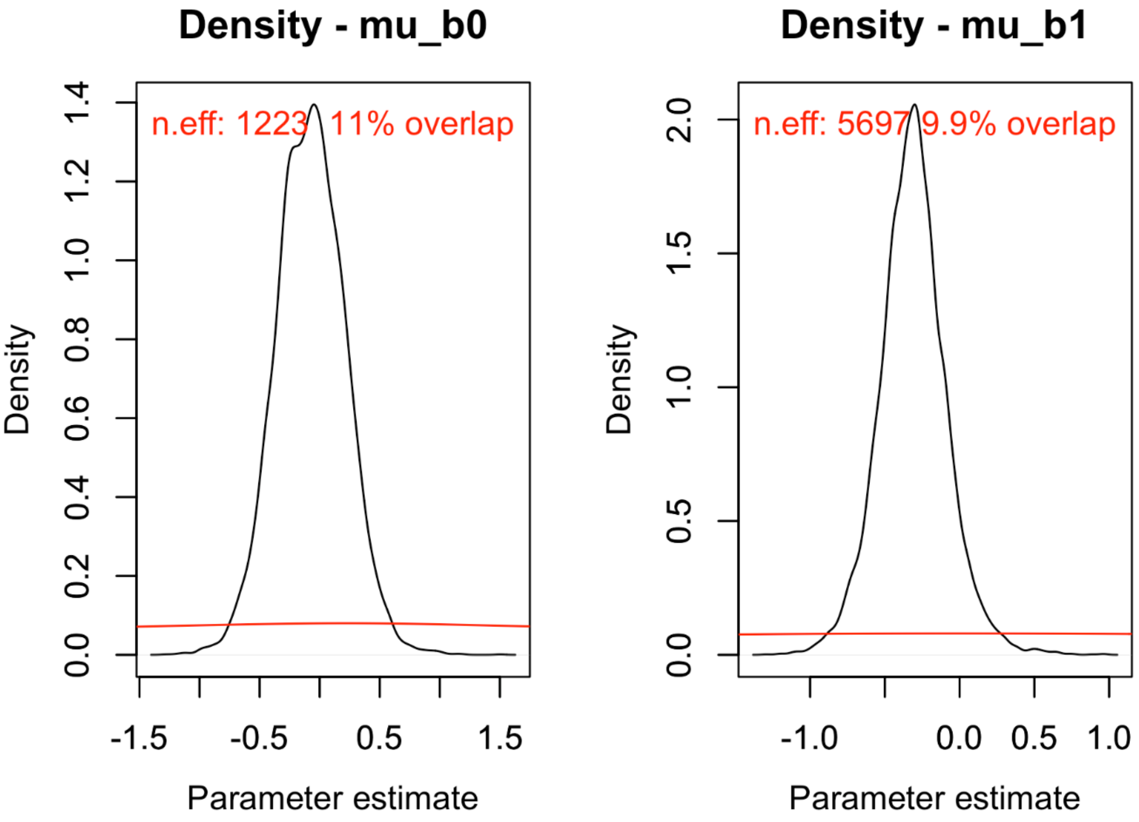

*Fig. S22. Posterior (black) and prior distribution (red) for the global parameters in the model for $T_{opt}$, including their % overlap and effective sample size (n.eff).*
